# Strong binding activity of few transcription factors is a major determinant of open chromatin

**DOI:** 10.1101/204743

**Authors:** Bei Wei, Arttu Jolma, Biswajyoti Sahu, Lukas M. Orre, Fan Zhong, Fangjie Zhu, Teemu Kivioja, Inderpreet Kaur Sur, Janne Lehtiö, Minna Taipale, Jussi Taipale

**Affiliations:** Division of Functional Genomics and Systems Biology, Department of Medical Biochemistry and Biophysics, and Department of Biosciences and Nutrition, Karolinska Institutet, SE 141 83, Stockholm, Sweden; Genome-Scale Biology Program, P.O. Box 63, FI-00014 University of Helsinki, Finland; Department of Oncology-Pathology, Science for Life Laboratory, Karolinska Institutet, SE 141 83, Stockholm, Sweden; Department of Biochemistry, University of Cambridge, United Kingdom

## Abstract

It is well established that transcription factors (TFs) play crucial roles in determining cell identity, and that a large fraction of all TFs are expressed in most cell types. In order to globally characterize activities of TFs in cells, we have developed a novel massively parallel protein activity assay, Active TF Identification (ATI) that measures DNA-binding activity of all TFs from any species or tissue type. In contrast to previous studies based on mRNA expression or protein abundance, we found that a set of TFs binding to only around ten distinct motifs display strong DNA-binding activity in any given cell or tissue type. Mass spectrometric identification of TFs revealed that within these highly active TFs, there were both housekeeping TFs, which were universally found in all cell types, and specific TFs, which were highly enriched in known factors that determine the fate of the analyzed tissue or cell type. The importance of a small subset of TFs for determining the overall accessible chromatin landscape of a cell suggests that gene regulatory logic may be simpler than what has previously been appreciated.

---

Transcription factors are proteins that read the gene regulatory information in DNA, and determine which genes are expressed. There have been many efforts to decipher the most important TFs in different cell types. Genetic analyses have revealed that cell identity is determined by a relatively few TFs that regulate each other, and bind often to specific regions of the genome together^1-3^. Cell identity can often be reprogrammed by exogenous expression of one to five TFs; for example, differentiated cells can be transformed into induced pluripotent stem (iPS) cells by expression of several different subsets of the TFs Oct4 (Pou5f1), Sox2, Klf4, c-Myc, and Esrrb^4-6^. These results indicate that the cellular regulatory system is hierarchical, and can be controlled by a relatively small subset of TFs.

Analyses based on gene expression profiling have, however, revealed that most tissues express hundreds of TFs^7^. Similarly, total proteomic analyses have indicated that tissues commonly express more than 40% of all ~ 1500 known human TF proteins^8^. Experiments based on chromatin immunoprecipitation followed by sequencing (ChIP-seq) have also suggested that a large number of TFs are active in individual cell lines^9,10^. Taken together, these results suggest that downstream of the master regulators, gene regulatory logic inside cells appears to be extremely complex, and that the cellular state could potentially be defined by a very large number of regulatory interactions. However, little information exists on which TFs have the strongest activities in a given cell type. This is because previous analyses have either only analyzed RNA or protein levels^7,8^, or measured individual TF activities using methods that cannot compare activity levels between TFs (e.g. ChIP-seq^9-11^).

To determine the relative activities of TFs in cells, we have in this work established a novel method, Active TF Identification (ATI). ATI is a massively parallel protein activity assay that can be used to determine absolute number of TF binding events from cell or tissue types from any organism. This information can then be used to derive relative DNA-binding activity of different TFs in the same sample. The word “activity” is used here in the same sense as in enzymology, where activity represents total enzyme activity (specific activity times molar amount). As ATI measures TF activity, and not occupancy at specific sites, it can be used to build models of TF binding based on biochemical principles that will contribute to our understanding of how DNA sequence determines when and where genes are expressed. In this work, we have used ATI to determine the relative distribution of biochemical TF activities in different cells, and the relationship between TF DNA-binding activities and overall chromatin architecture. We found that only few TFs display strong DNA binding activity in any cell or tissue type from all tested organisms. The strongly active TFs can be used to predict transcript start positions in yeast, and chromatin accessibility in mammalian cells. Our findings indicate that a small core set of TFs display much stronger DNA-binding activity than other TFs in all tested organisms and cell and tissue types. The TFs that have strong DNA binding activity play an important role in setting up the overall chromatin landscape, and in helping to open chromatin for TFs that display weaker DNA binding activity. Compared to a system where all TFs can cooperate with all other TFs, the resulting hierarchical organization leads to a greatly simplified cellular gene regulatory network.

## Results

### *De novo* identification of motifs bound by TFs from cell extracts

In ATI, a library of double-stranded oligonucleotides containing a 40 bp random sequence is incubated with a nuclear extract from different cell or tissue types. The oligonucleotides bound by TFs are then separated from the unbound fraction by electrophoretic mobility shift assay (EMSA, **Fig. 1a**). The bound DNA fragments are eluted from the gel and amplified by PCR, and the entire process is repeated three more times. Comparison of millions of sequences derived from the input and the selected libraries then allows identification of enriched binding motifs that correspond to the TFs present in the nuclear extract. Given that the binding motifs identified are relatively short compared to the 40 bp random sequence, the sequence flanking the motif can also be used as a unique molecular identifier^12^, allowing absolute quantification of the number of proteins bound to each type of motif.

**Figure 1.**
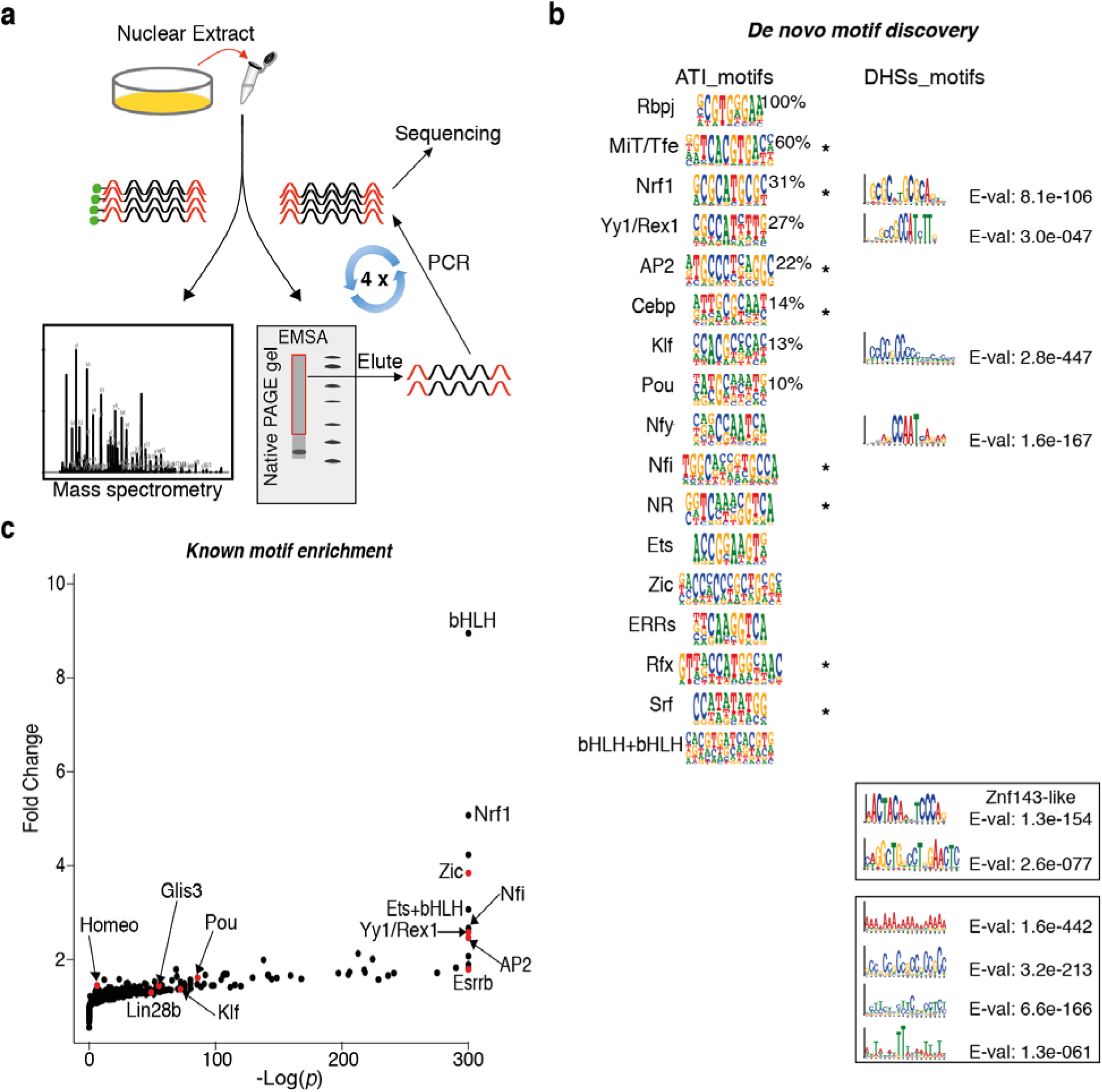
Active transcription factor identification (ATI) assay. **a,** General description of the process. After incubation of proteins extracted from cells or tissues with double-stranded DNA oligonucleotides containing 40 bp random sequences, oligos bound to the proteins are separated from unbound DNA by native PAGE gel purification (EMSA; right) and amplified by PCR. The process is repeated three more times, resulting in enriched DNA oligonucleotide pool that reflects activities of the TF proteins in the cell lysate. The ATI assay consists of two parts: motif analysis and MS identification of the TFs. The sequencing data for original and enriched DNA oligos is used for motif analysis (**b** and **c**); the MS identification of the TFs was carried out by capturing proteins from the nuclear extract by a biotinylated oligonucleotide pool followed by mass spectrometry (left). **b,** Logos of motifs discovered using *De novo* motif discovery from ~ 10 million ATI sequence reads (“ATI_motifs”, based on Autoseed program^13^) and from top ~ 1000 DHS regions (“DHS_motifs”, based on MEME) in mouse ES cells. In the category of “ATI_motifs”, names of TFs are based on motif similarity to known motifs (see **Supplementary Fig. 1a** for details). The motifs are quantified with background corrected absolute molecular counts^12^ at cycle 4; the highest count is normalized to 100%. Motifs with counts more than 10% of the maximum are considered “strong” motifs; for these motifs, the relative molecular counts are indicated in the right corner of corresponding motif. Asterisk indicates homo-dimeric motifs bound by TFs from the same family. In the category of “DHS_motifs”, top 10 motifs with the lowest E values are shown: four motifs that are similar with related ATI derived motifs are displayed at the same rows of the related ATI motifs; two non-repetitive motifs including the Znf-143 like motif and four repetitive motifs are displayed in the two separate boxes. **c,** Known motif enrichment analysis of ATI data from mouse ES cells. Known motifs are matched to the unselected and ATI enriched DNA sequences, and *p*-value (x-axis, log scale; due to the precision of calculation, many *p*-values are set to a minimum of 10^−300^) and fold change (y-axis) are calculated for each known motif. The enrichment of motif matches is correlated with DNA binding activities of different TFs or TF dimers in the nucleus. Red dots represent TFs that have been previously known to be important for maintaining pluripotency in mouse ES cells.

As an initial test, we performed ATI using nuclear extract from mouse ES (mES) cells cultured without feeder cells. *De novo* motif discovery using the “Autoseed” program^13^ revealed motifs characteristic of TF families such as Nfi, Rfx, Klf and Pou, and of subfamily specific motifs for MiT/Tfe basic helix loop helix proteins, class I ETS factors, Zic zinc fingers and ERR type nuclear receptors (**Fig. 1b**; for motif similarity, see **Supplementary Fig. 1a**). Many motifs were similar to motifs bound by known lineage-determining factors for ES cells^14-18^, such as Klf4, Pou5f1 (Oct4), Zic3 and Esrrb, suggesting that the ATI assay is able to detect the DNA-binding activity of such TFs. Some, but not all, ATI motifs were also detected by MEME motif mining of DNase I hypersensitive sites from ES cells, suggesting that ATI can detect also activities of TFs that contribute to chromatin accessibility (**Fig. 1b**). Analysis of the amount of DNA recovered from the gel, and the absolute number of motifs recovered during sequencing revealed that TFs from one nucleus bound approximately 5 × 10^6^ DNA ligands, of which 3 × 10^5^ represented specific binding events with clearly identifiable motif. This estimate is broadly consistent with earlier estimate of TF abundance^19^.

### Most recovered motifs are previously known

Although the *de novo* motif discovery using Autoseed is relatively sensitive, and can identify motifs that represent ~ 5 ppm of all sequences, it cannot identify very rare events. To detect such events, we also analyzed the enrichment of known motif matches during the ATI process (**Fig. 1c** and **Supplementary Table 2**). This method yielded similar results to the *de novo* method, but also identified enrichment of motif matches for additional TFs, including homeodomain motifs that are similar to motif bound by the pluripotency regulator Nanog, indicating that it has higher sensitivity than the *de novo* motif discovery method. The relatively low enrichment of the homeodomain motif is consistent with its relatively low abundance of Nanog in the mass spectrometry analyses (**Supplementary Table 3**). However, even using known motif enrichment, it is hard to unambiguously assign a weak activity to a specific TF. This is because it is difficult to determine whether enrichment of the motif matches that is too weak to be detected by *de novo* methods represents specific enrichment of the tested motif, or is a consequence of stronger enrichment of a related motif of another TF.

It is interesting to note that all of the motifs recovered from ES cells were known prior to this study, suggesting that few novel strong TF activities remain to be discovered. Among those strong motifs, there are both monomeric motifs bound by one single TF and dimeric motifs bound by dimers formed by two TFs from the same structural family (**Figure 1b**). However, some motifs representing dimers formed on DNA^20^ such as the well-known Sox2/Pou5f1 motif were absent from the data. This result suggested that the ATI method might be biased against such DNA-dependent dimers, as far larger number of monomeric than dimeric sites exist in random sequences (**Supplementary Fig. 1b**). To address this, we performed two additional experiments using a synthetic library consisting of known motifs, and a library derived from mouse genomic sequences. These targeted analyses were more sensitive than the method based on random library, and resulted in identification of many additional motifs, including that of CTCF (**Supplementary Table 4**). However, analysis of the data did not reveal DNA-dependent dimeric motifs, suggesting that the activity of DNA-dependent TF dimers is lower than that of the corresponding monomeric TFs. This is likely because specific formation of DNA-dependent dimers requires that the binding activity of at least one of the TFs is relatively low, as its concentration must be below the individual Kds towards the motif and the partner, but above the Kd towards the combination of the dimer motif and the TF partner. Due to the fact that proteins are more concentrated in the nucleus than in the nuclear extract, such dimers are difficult to detect using ATI.

### Identification of specific transcription factors by mass spectrometry

The ATI assay can be used to identify active motifs, but analysis of motif activity alone cannot in most cases identify the specific TF that is active in the tissue or cell types, due to the fact that many related proteins can bind to the same sequence motif. To address this, we captured DNA-binding proteins from the nuclear extract in mouse ES cells using a biotinylated version of the control and enriched ATI DNA ligand libraries. After washing and elution, we identified proteins bound to the ATI ligands using mass spectrometry (see **Methods** for details). This analysis revealed the TF proteins that bound to the ligand. In most cases, a motif could be assigned to a specific protein or a group of paralogous proteins (**Supplementary Table 1**).

### Highly active TFs can be classified to common, shared and specific classes

We next applied ATI to identification of TFs active in mouse ES cells and four adult mouse tissues, including heart, spleen, brain and liver. *De novo* motif discovery followed by motif match counting revealed that limited sets of TFs were highly active in different tissue types (**Fig. 2a** and **Supplementary Table 5**). In all tissues tested, only few TFs (two to seven) displayed activities that were more than 10% of that of the most active TF. The identified motifs could be broadly classified into three groups: common, shared and specific. Five common motifs were found in all cell and tissue types tested. They represented an extended E-box site (gGTCACGTGACc) bound by the MiT/TFE family of basic helix-loop helix (bHLH) TFs, a GGTCAaaGGTCA motif bound by a subfamily of nuclear receptors (NR), and canonical sites bound by NFI, NRF and bZIP (Creb) family of TFs (**Fig. 2a**; see **Supplementary Fig. 2a** for comparison to known motifs). Even within this common set, there were large differences in quantitative TF binding activities between the cell types, suggesting that the relative activities of the common TFs may contribute to cell lineage determination (**Fig. 2b** and **Supplementary Fig. 2b**).

**Figure 2.**
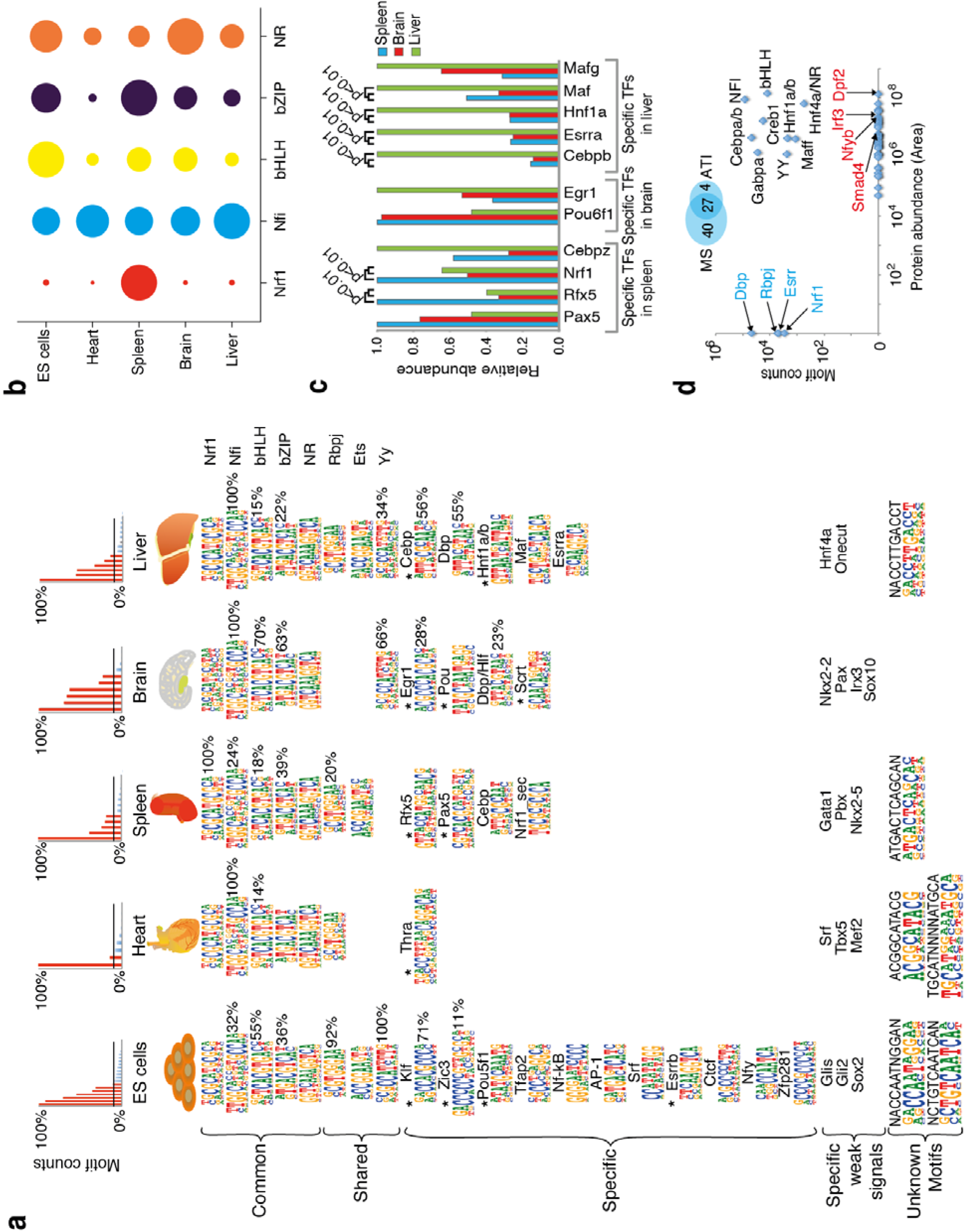
Deciphering most strongly active TFs in different cell types. **a,** Only few TFs are strongly active in mouse ES cells and tissues. Top: Histogram shows background corrected absolute molecular counts^12^ (y-axis, Motif counts) of all discovered motifs at cycle 4; the highest count is normalized to 100%. Counts more than 10% of the maximum are indicated by red bars, and the corresponding motifs are considered as “strong” motifs; the relative activities of them are shown on the right corner of the corresponding sequence logos. Bottom: Sequence logos and the corresponding TFs identified by *de novo* motif discovery from the indicated tissues are shown (see **Supplementary Table 8** for PWM models of all logos). Five motifs are found in all samples (common), whereas three motifs are found in many but not all samples (shared). Many motifs are also found in only one or two tissue or cell types (specific), including several factors known to contribute to lineage determination in the analyzed tissues (indicated by an asterisk). Examples of TFs known to be important for the specific tissues whose motifs were only identified using the known motif discovery pipeline are also indicated (specific weak signals; see **Supplementary Table 6** for details). Some unknown motifs were also identified. The names of the TFs are based on the motifs (see **Supplementary Fig. 2a** and **2c**). In cases where multiple TFs are known to bind the same motif, the motif is assigned to the specific TF based on mRNA expression level and functional data from previous studies (see “References” in **Supplementary Figure 2c**). **b,** Quantitative levels of DNA binding activities of the common TFs vary between the tissues. The areas of circles indicate the activities of the five common motifs in the indicated tissues, based on increase of absolute molecular counts^12^ of each motif in the sequencing data from the original library (cycle 0) to the last cycle (cycle 4). The activities of each TF are normalized by setting its highest activity in any of the tissues as 1. **c,** Relative protein levels of the ‘specific’ TFs as measured by TMT-labeled mass spectrometry from triplicate samples correlate partially with the corresponding TF activities measured by ATI motif analysis. The relative abundance of each TF protein is normalized by setting the highest abundance as 1. *p*-values for significant differences are also given. The abundance of Egr1 (**asterisk**) differs between replicate samples from different mice based on both types of MS analyses, suggesting that its abundance varies naturally in mouse liver (see **Supplementary Table 9 and 10**). **d,** Label free mass spectrometry detects many TFs that are strongly active (black font; y-axis, ATI motif counts), but also reveals high protein abundance (x-axis) of TFs that are not strongly active in mouse liver, including Dpf2, Irf3, Nfyb and Smad4 (red font). The MS analysis also fails to detect some TFs whose motifs are recovered by ATI (blue font; Rbpj, Dbp, Esrr, Nrf1). Note that the same motif detected in ATI could be correlated with several different TFs detected in the mass spectrometry.

In addition to the common TF motifs, we also found three motifs, corresponding to Rbpj^21^, class I Ets family TFs^22^ and Yy1/2^9^ that were shared by more than two different tissue types, suggesting that members of these families of TFs have important roles in many different contexts (**Fig. 2a**; see **Supplementary Fig. 2a** for comparison to known motifs). In contrast, there were some other motifs specific for only one or two tissue types (**Figure 2a**; see **Supplementary Fig. 2c** for comparison to known motifs); some of their corresponding TFs have been previously shown to be crucial for the particular cell identities. For example, the binding motif of nuclear receptor Thra that is important for heart function^23^ was found only in the heart, whereas the Rfx and Pax motifs were found only in the spleen, where it is known that members of these families such as Pax5 and Rfx5, respectively, contribute to development^24,25^ and MHC class II expression^26^ of B-cells. In the brain, we detected motifs for Egr, Scrt and Pou2, Pou3 (Brn2) and Pou6 (Brn5) family proteins that are specifically expressed and play important roles in the brain^27–30^. In the liver, motifs bound by Cebp family TFs, Hnf1a/b and PAR-domain bZIP TF Dbp/Tef/Hlf were specifically enriched, and these TFs have been verified to be crucial for liver functionality as well as circadian control of metabolism^31–37^. Moreover, it has been shown that Hnf1a/b and Cebpa together with other factors can be used to reprogram fibroblasts into induced hepatocytes^38,39^, indicating the significance of these factors for hepatocyte identity.

### Many TFs that determine the cell fate are strongly active in cells

We next analyzed the relative enrichment of known TF motifs across the tissues using known motif enrichment analysis. This analysis confirmed the enrichment of the *de novo* discovered motifs, and revealed additional TFs whose motifs were specifically enriched in the different cell types or tissues (**Fig. 2a** and **Supplementary Table 6**). For instance, from the mES cells we detected motifs for additional pluripotency factors such as Glis2, albeit with a relatively low enrichment. From adult mouse liver, we also detected specific motifs for TFs such as Onecut and Hnf4a. Taken together, ATI analysis of mouse tissues revealed that in addition to five common TF activities, each tissue displayed strong activity of key regulators for the respective cell identities.

To test if TFs detected by ATI can induce transdifferentiation of somatic cells to other cell identities, we transduced human fibroblasts with a combination of nine specific TFs identified by ATI from adult mouse liver, and investigated the morphology of the cells and expression of the liver specific marker albumin after two weeks of culture. The fibroblasts were converted to induced hepatocytes (iHeps) at an efficiency that was similar to the most efficient previously described protocol (**Supplementary Fig. 3**), indicating that ATI can identify all factors necessary for transdifferentiation of mammalian cells.

We also detected enrichment of some unknown motifs (**Fig. 2a**, bottom), which we could not assign to a known TF based on current knowledge (HT-SELEX motifs, CIS-BP, TOMTOM^9,13,40,41^). Overall, we recovered 35 motifs, of which only 6 (17%) were unknown, indicating that specificities for most TFs that display strong activity in the tested tissue types have already been determined.

### TF activity is not explained by protein abundance alone

To identify the TFs, we carried out mass spectrometry (MS) analyses in three adult mouse tissues: spleen, brain and liver. This analysis was performed using HiRIEF-LC-MS^42^ with relative quantification between samples using isobaric tags (TMT). Comparison of the MS data and ATI motif analysis result revealed that most of the TF proteins whose motifs were specific for one of the tissues were more abundant in that tissue compared to the other two tissues (**Fig. 2c**). We also performed label-free MS analysis to estimate the protein levels within the samples. In this analysis, we detected some highly abundant TFs that were not detected in the ATI assay; for example, no motif was discovered for Dpf2 that is involved in apoptosis^43^. Similarly, Smad proteins were abundant in all of the tissues, but their signature motif was not detected in any of the cases (**Fig. 2d** and **Supplementary Table 7**). This is consistent with low DNA binding activity of Smads^44^, and the fact that Smad protein activity depends on posttranslational modifications induced by stimulation of cells by TGF-β superfamily ligands. We also found that interferon regulatory factors (IRFs) were abundant at protein level, yet not detected as active by ATI. In addition, the MS analysis indicated that some subunits of the multimeric TFs such as NFY were present at high levels, yet the TFs were not strongly active, consistent with lower level of abundance of the other subunits. Of the 67 TFs that were detected by mass spectrometry in liver, 40% were known to bind to an ATI identified motif, 12% represented a known obligate heteromer or ligand-regulated TF, 33% were classified as TFs but do not have a known motif, and 15% had a known motif but were present at a relatively low abundance (**Supplementary Table 7**). In summary, the ATI assay revealed that the total DNA binding activities of TFs in cells cannot be simply determined by measuring RNA or protein abundance alone, as many TFs either have low specific activity, or are only activated under specific circumstances.

### Strength of DNA-binding activities change during cell differentiation

To test if ATI can detect changes in TF activities that are induced during cell differentiation, we induced differentiation of ES cells towards neural and mesodermal lineages using standard conditions^45-47^ (**Fig. 3a**; see **Methods** for details). This analysis revealed that ATI was able to detect several known quantitative changes in TF binding activities that accompany the neural differentiation process (**Fig. 3b** and **c**). For example, the activities of the pluripotency factors Glis and Zic were decreased, whereas the activity of Rfx and Pax factors that are known to contribute to neural differentiation was increased. Similarly, the activities of Glis and Zic factors decreased after induction of mesodermal differentiation, whereas the activity of the known mesodermal factor AP2 increased dramatically (**Fig. 3b** and **d**). The activation of Smad proteins by BMP4 and Activin A that were used to induce mesodermal differentiation was not detected, potentially due to the fact that Smad proteins bind DNA only weakly (Kd ≈ 1 × 10^−7^ M), and often act together with other TFs^44^. In contrast, ATI robustly detected the activation of retinoic acid receptor by the neural inducer retinoic acid (**Fig. 3b** and **c**), indicating that some ligand-gated TFs can be detected by ATI.

**Figure 3.**
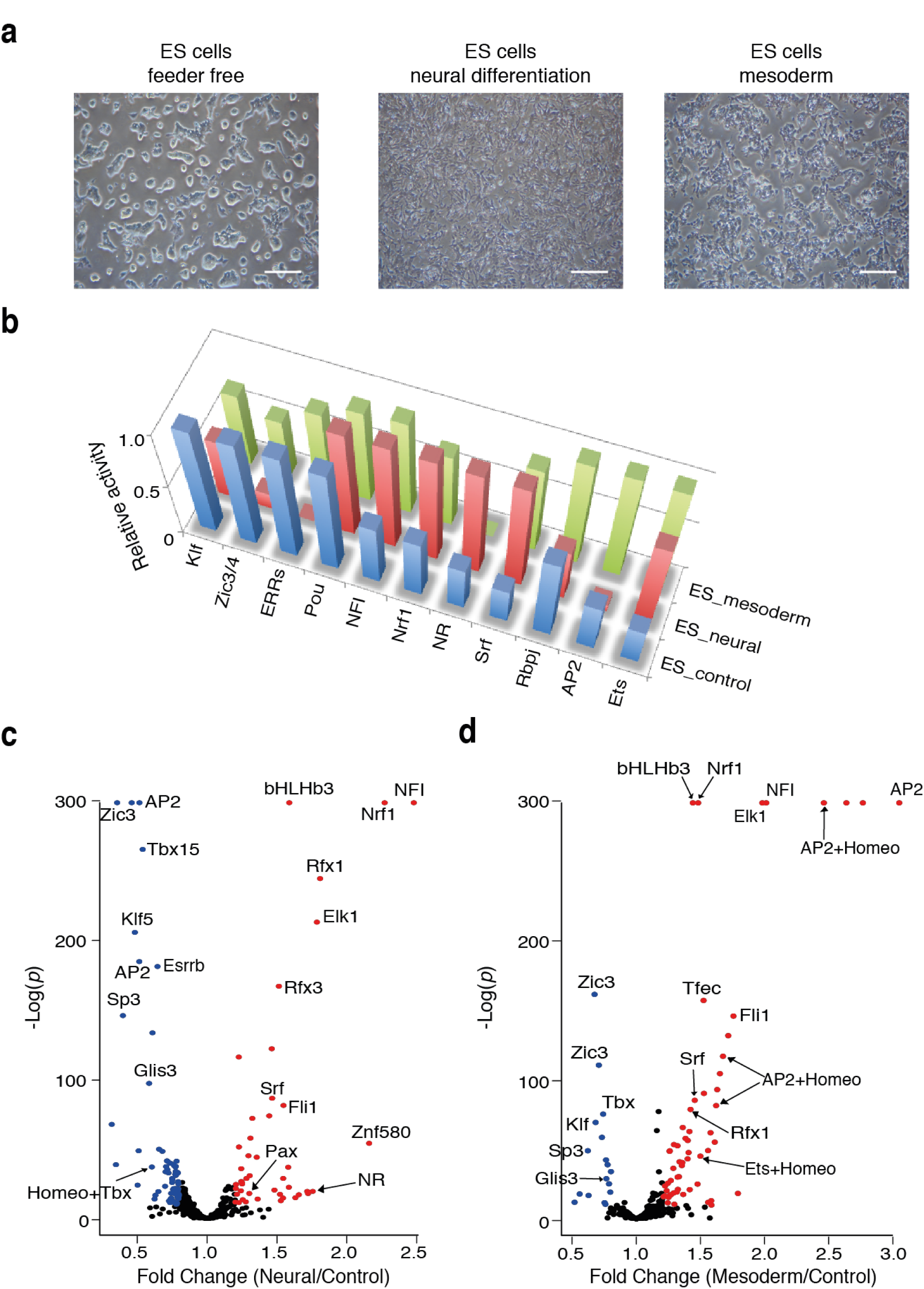
ATI analysis of TF activities in differentiating ES cells. **a,** Morphology of control mouse ES cells (left), and ES cells induced to differentiate towards neural (middle) and mesodermal (right) lineages two days after induction, scale bar = 400 μm. **b,** Comparison of the motifs detected by *de novo* motif discovery method in the ES cells and differentiated ES cells. Bars indicate the relative activities of the indicated TFs based on increase ^12^ of the absolute molecular counts of each motif between the first cycle and the fourth cycle of ATI. The activities of each TF are normalized by setting the highest activity in any of the three conditions to 1. **c,** Comparison of motif enrichment between the neural differentiated ES cells and the control ES cells. y-axis: *p*-value (log scale; due to the precision of calculation, many *p*-values are set to a minimum of 1 × 10^−300^). x-axis: fold change. The motifs with *p*-value less than 1 × 10^−10^ and change of more than +20 % or −20% are indicated by red or blue colors, respectively, with names of some motifs indicated. **d,** Comparison of motif enrichment between the mesodermal differentiated ES cells and the control ES cells. y-axis: *p*-value (log scale; due to the precision of calculation, many *p*-values are set to a minimum of 1 × 10^−300^). x-axis: fold change.

### ATI reveals conservation of TF binding activities between distant species

One of the important advantages of the ATI assay is that it can be performed using any type of protein extract from any species. To analyze how similar active TFs are between organisms, we carried out ATI experiments using nuclear extracts from the fruit fly *D. melanogaster* S2 cells, and the yeasts *S. cerevisiae* and *S. pombe*. Analysis of the sequencing data using the *de novo* motif discovery method indicated that ATI could identify the most active TFs in all of these species. Several of the recovered motifs matched known motifs from the respective species^41^. From the study of the yeast *S. cerevisiae*, we detected Abf1, Rap1 and Reb1. Strikingly, of six motifs (five common motifs and one shared motif for Rbpj) that were common to almost all mouse tissues and cell lines, two TFs, Rbpj/Cbf11 and the bHLH factor Tfe/Cbf1, were highly active in the yeast *S. pombe,* and two TFs Tfe/Cbf1 and the bZIP factor Creb/CST6 were highly active in *S. cerevisiae*. Although there are only 23 TF families, the number of distinct motifs is much larger; for example in humans, there are more than 300 different binding motifs^48^, and at least 30 distinctly different motifs for bHLH factors, and 10 for bZIP factors^9,48^. Thus, the fact that the same motifs were highly active in distant species suggests that the dominant mechanisms of transcriptional regulation may have been conserved during the evolution of eukaryotes. In the different species, we also found many novel motifs, which we could not assign to a known TF based on existing TF specificity databases, suggesting that the binding specificity landscape of these species is not characterized as well as that of mammals (**Supplementary Fig. 4**).

### Identification of master TFs that determine the transcriptional state of yeast

One concern with the ATI assay is that it could identify specific TFs that bind strongly under the *in vitro* conditions used in the assay. To independently validate the assay, we analyzed whether the most active DNA-binding TFs identified by the method were also the most active *in vivo.* We first compared the ATI results from the yeast *S. cerevisiae* with the known four TF motifs that determine the yeast transcriptome^49^. Out of four motifs that are required to build a model describing yeast transcript start positions, ATI identified three Abf1, Rap1 and Reb1, and in addition recovered novel CG rich motifs that are related to the fourth motif used in the study (Rsc3; **Fig. 4a**). This result indicates that ATI can identify a complete set of TFs whose activity is crucial for determining the transcriptional state of yeast.

**Figure 4.**
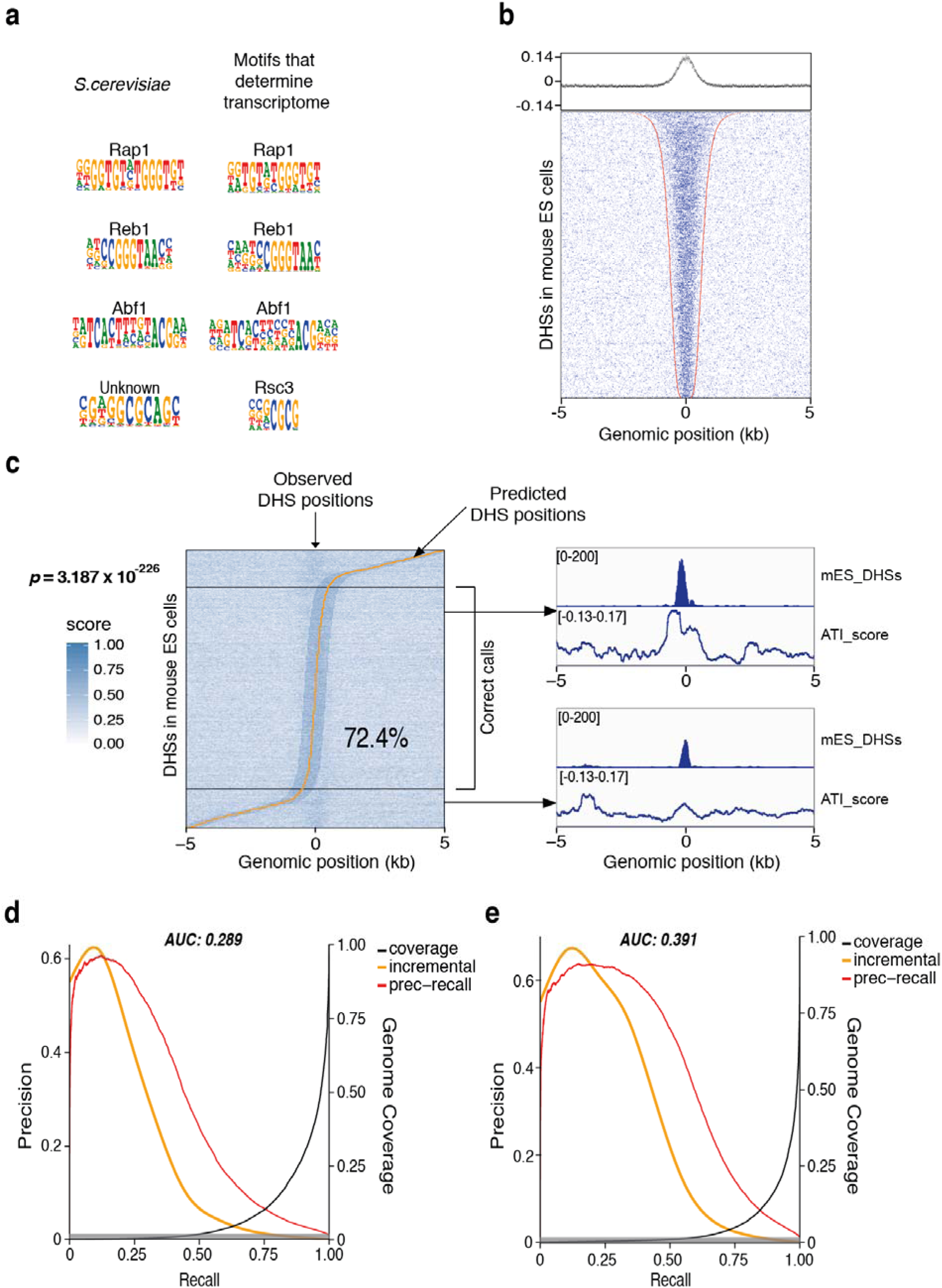
Strongly active TFs explain key features of transcription in yeast and in mouse cells. **a,** Comparison of motifs detected in ATI assay (*S. cerevisiae*) and the four motifs that can be used to computationally identify yeast transcript start positions^49^. **b,** ATI enriched 10-mers from mouse ES cells are also enriched in DNase I hypersensitive sites from ES cells. In the dot plot, each row indicates one DHS region from the ES cells that is flanked with genomic sequences. Red dots indicate the boundaries of the DHS regions, blue dots indicate positions of top 2000 ATI-enriched 10-mers out of all 4^10^ (~ 1 million) 10-mers. The graph on top shows the average of scores for each 10-mer at each position across the rows. **c,** Prediction of ES cell DHS regions by using the 10-mer data from the ATI assay. DHSs are sorted by position of the prediction call (yellow line). Black horizontal lines separate accurate DHS calls (middle) from calls more than 500 bp off the known DHS center that is located at the x-axis position 0 in all cases. The fraction of predictions within ± 500 bp of the center and the corresponding *p*-value for null model where position calls are randomly distributed are also indicated. Two example tracks for the DHS and ATI signals are also shown. **d-e,** Comparison between the genome-wide predictions of the ES cell DHS regions using 10-mer data from the ATI assay (**d**) and DHSs themselves (**e**). Precision-recall curves (red lines, cumulative precision) indicate that the ATI 10-mers can be used to predict the DHS positions (AUC 0.29), and that the performance of the ATI predictor is relatively close to a predictor that uses 10-mer data from the DHSs themselves (AUC 0.39). Yellow and black lines, respectively, show smoothed incremental precision and fraction of the genome selected at the indicated recall level. Gray shading indicates fraction of true DHSs in the genome (0.9%).

### TF binding activities contribute to open chromatin landscape of mammalian cells

To determine if ATI also confers information on the mammalian chromatin landscape, we compared the ATI data with DNase I hypersensitive sites (DHS) from the mouse ENCODE project^50^. This analysis revealed that the top 2000 10-mers detected by ATI in mouse ES cells, heart, spleen, brain and liver were strongly enriched in the ~ 5000 most significant DHS regions from the respective tissues. As expected, the strongest enrichment was seen in ES cells, which are more homogenous than tissues containing multiple different cell types (**Fig. 4b** and **Supplementary Fig. 5**). Analysis of 10-mers enriched in ES cell DHSs and ATI revealed that there were many 10-mers that were enriched in both, and that all of these 10-mers were related to the ATI motifs (**Supplementary Fig. 6**). Consistent with the enrichment of ATI 10-mers in DHSs, ATI also enriched many DHS sequences from a mouse genomic library (**Supplementary Fig. 7**). However, some 10-mers were only enriched in DHSs (**Supplementary Fig. 6**); these included many repetitive CG rich sequences that enrich in gene regulatory elements due to the fact that methylated C is prone to mutation, and the low CpG methylation rate of regulatory elements protects these sequences from this mutational process^51,52^. As DHSs represent gene regulatory elements, they are also expected to be enriched with both motifs that contribute to the opening of the chromatin, and motifs that are involved in downstream activities such as transactivation or repression of RNA polymerase II. Consistently, *de novo* motif discovery analysis of DHSs revealed some motifs that were not enriched by ATI. These included a motif similar to that of Znf-143 (**Fig. 1b**); this motif has been reported to contribute to interactions between promoters and distal regulatory elements^53^. Interestingly, *de novo* motif discovery analysis of both ATI and DHS data failed to detect DNA-dependent dimer signals, suggesting that either such dimers are not commonly strongly active, or that many different dimers contribute to opening of chromatin at different DHSs.

We further hypothesized that if ATI accurately represented TF binding activities in cells and revealed subsequences that bound strongly to TFs also *in vivo*, it would be possible to predict the DHS regions using the ATI enriched subsequences. It has been previously shown that DHSs can be predicted based on sequence features from different types of experimental data (e.g. DNase-seq data^54^ or ChIP-seq data^55^). It is also well established that DHSs and TF binding clusters are enriched with matches to biochemically obtained TF motifs^56,57^, and that they overlap with *in silico* predicted clusters called based on TF motif matches only. However, in our recent study, only ~ 30% of TF binding clusters could be predicted based on monomeric TF binding models^57^ suggesting that additional unknown determinants affect TF binding to DNA inside cells. To determine if ATI improves on the predictions, we developed a predictor based on the enrichment rank of all 10-mers in ATI. This analysis revealed that more than 70% of the DHSs could be predicted by the 10-mers derived solely from ATI (**Fig. 4c**; 10% expected by random, *p* < 3.2 × 10^−226^; winflat^58^), indicating that ATI derived 10-mer enrichment more accurately represented TF activity in cells compared to any other presently available information. In prediction of the DHSs genome-wide, ATI was nearly as effective as using 10-mers from the DHSs themselves (**Fig. 4d, e**), indicating that the ATI data contains substantial fraction of the motif information included in the DHSs, despite the fact that the DHSs are expected to contain additional motif features that relate to their functionality in gene regulation and not to their open chromatin status. Consistently, analysis of DHSs that were hard to predict using ATI 10-mers but could be predicted using DHS 10-mers again revealed the Znf-143 like motif (**Supplementary Fig. 8d**). In contrast, DHSs that were hard to predict using both types of 10-mers did not contain enriched sequence motifs (**Supplementary Fig. 8d**), suggesting that their DNase I hypersensitivity may be caused by longer sequence features such as those affecting intrinsic nucleosome affinity^59^. In summary, both the yeast transcriptome model and the DHSs data verify that the TFs found using ATI are active in the cell types and tissues analyzed, and contribute to the positioning of TSSs in yeast, and accessibility of chromatin in mouse cells.

## Discussion

It is not possible to determine the binding activities of TFs in different types of cells based solely on RNA expression or protein abundance data. This is because different TFs have different binding specificities and affinities to DNA. Moreover, the binding activity of TFs is often regulated at the level of posttranslational modification, protein-protein interaction^60^, and nuclear localization. For this reason, we have in this work developed the ATI assay that directly measures the DNA-binding activities of all TFs in cell extracts. The ATI method is widely applicable, as it can be applied to any species and tissue, and can detect changes in TF activity in response to any perturbation. This method is, to our knowledge, the first massively parallel protein activity assay that can simultaneously measure a large number of functionally similar protein activities. The method can be used to determine the overall distribution of TF activities in cells, and the change in the activities in response to stimuli. In addition, comparison of TF activities between different cell types yields useful hypotheses about key transcription factors that determine the identity of the cell types. Based on independent verification of the results using mass spectrometry and prediction of functional features, the method appears to be able to capture the majority of strong DNA-binding activities in cells. However, the sensitivity of ATI in detecting accessory DNA-binding factors and proteins that bind to DNA weakly such as the Smad proteins may be low. Further studies that miniaturize and standardize the process are expected to further improve the sensitivity and accuracy of this widely applicable method.

Analysis of active TFs in different species revealed a potential conservation of strongly active TFs between distant species. In addition, we found that in higher organisms, relatively few TFs display strong DNA binding activity in any given tissue type. From different mouse tissues, we detected five TF classes that were commonly and strongly active. In addition, we found many TFs that were strongly active only in specific tissues; these TFs had known roles in cell lineage determination. Differentiation of ES cells towards neural and mesodermal lineages was associated with rapid changes in TF activities. In addition, we were able to transdifferentiate human fibroblasts towards hepatocytes by expression of a novel, ATI-identified combination of liver-specific TFs. These results suggest that strongly active TFs have an important and active role in cell differentiation and cell fate determination.

We also found that using the ATI-derived binding information we were able to predict positions of open chromatin in mouse ES cells far more accurately than what has previously been possible, indicating that lack of knowledge of TF DNA-binding activity levels was a major unknown factor that hindered previous computational predictions of regulatory elements. These results suggest that strongly active TFs have a major role in setting up overall open chromatin landscape of cells. However, our results cannot be interpreted to mean that open chromatin would result exclusively from the action of TFs with strong binding activity, or that binding motifs of strongly bound TFs would be sufficient to open closed chromatin states characterized by presence of HP1, histone H1 or repressive chromatin modifications^61^. It is well known that some TFs can directly or indirectly recruit enzymes that remodel or modify nucleosomes to generate open chromatin and/or derepress closed chromatin states^62,63^. It should also be noted that large number of binding sites for less active TFs, or site(s) for combinations of cooperatively bound TFs can also bind with sufficient energy to displace or compete out nucleosomes. These mechanisms can open a subset of DHSs, but are unlikely to be the predominant way to open chromatin, as if that was the case, the binding motifs for the cooperative or weak binders would have been detected based on the *de novo* motif mining of the DHS regions.

Our results indicate that the few TFs that have strong DNA binding activities in a cell play a major role in setting its overall gene regulatory architecture. On the other hand, ChIP-seq analyses have clearly shown that a large number of TFs can bind open chromatin regions in cells^9,10^. These observations are consistent with a model where the TFs that are strongly active in DNA binding set up the overall chromatin state of cells, and that the ability of TFs with weaker DNA binding activity to regulate their target genes is conditional on this chromatin state.

Compared with a complex model where different TFs are equally active and can collaborate with each other, such a hierarchical gene regulatory model is far more simple, and can explain the fact that hierarchical gene expression patterns are commonly observed in analyses of real biological systems (**Figure 5**). Interestingly, this model also provides a very simple combinatorial gene regulation system. If the TF that has strong DNA-binding activity lacks a strong transactivation or repression domain, it will require a partner that has such a domain. This co-operating factor may not, in turn, be able to bind DNA strongly to open chromatin alone, and therefore will require the strong DNA binder. It should be noted that different types of activation domains will also further contribute to such combinatorial regulation^64^, increasing the number of cooperation partners to three or more.

**Figure 5.**
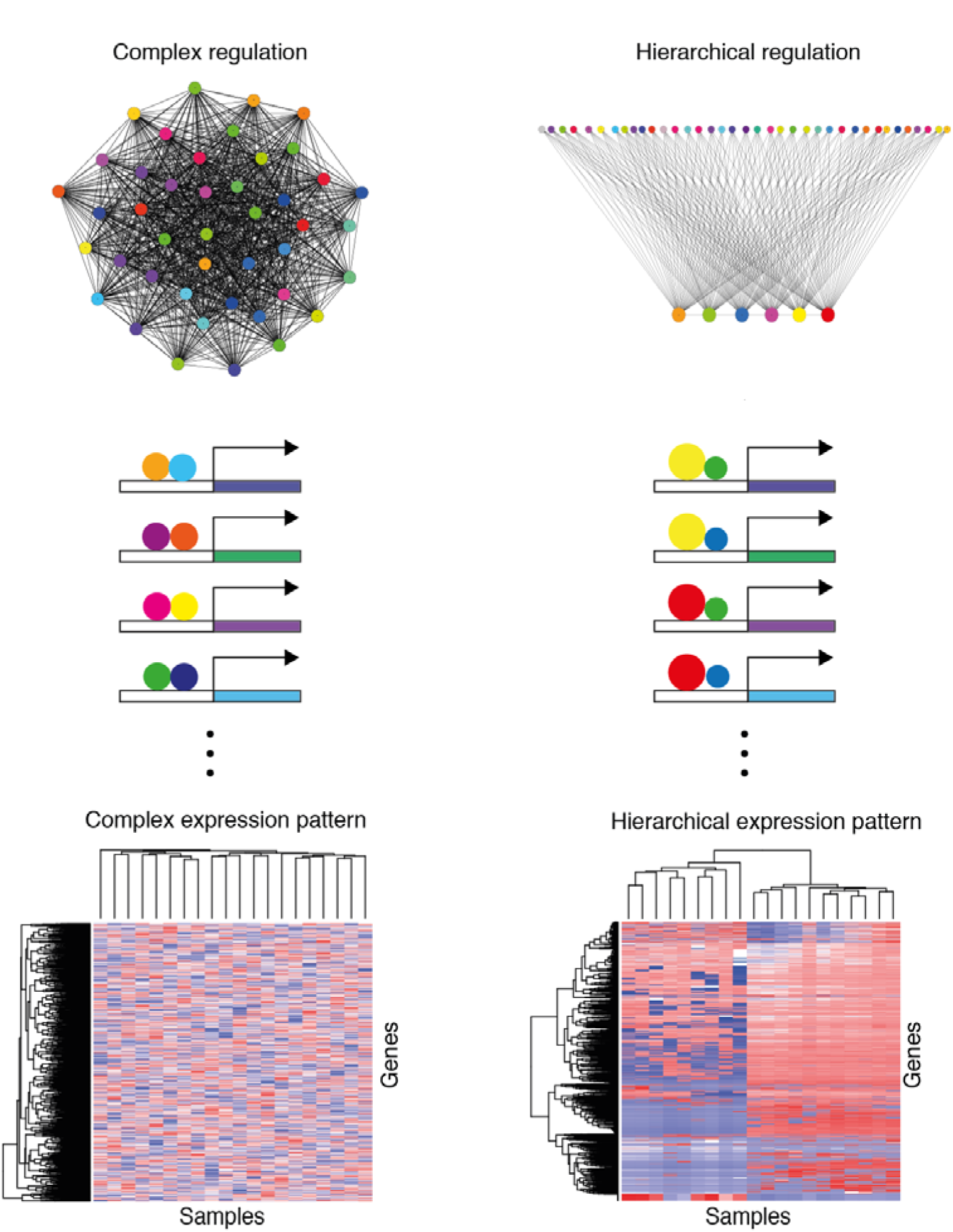
Hierarchical gene regulatory network leads to a Hierarchical gene expression profile. Compared with a complex gene regulatory network formed from equally active TFs, the hierarchical gene regulatory network formed using TFs that are either strongly (large circles) or weakly (small circles) active is simpler, and yields a gene expression pattern that is similar to the hierarchical gene expression patterns observed in real biological samples. The heatmap for complex expression pattern is generated from an artificial random matrix containing 20 “samples” and 2,000 “genes”; the heatmap for hierarchical expression pattern is generated from RNA-seq data from 20 different cell and tissues samples in the ENCODE project^50^. In this model, the genes clustered into each group are regulated by the same dominant TFs in collaboration with different weak TFs, and clustered tissue samples share some dominant TFs.

The biochemical activity-based distinction between TFs that we identified here is related but not identical to the concept of “pioneer” transcription factors that are able to bind to nucleosomal DNA^65^. Ability to compete against nucleosomes could either be simply due to mass action, or due to a specific ability of some TFs to bind to nucleosomal DNA with fast kinetics and to recruit nucleosome remodeling enzymes^62,63^. In bioinformatic studies that identify factors occupying most of their consensus motifs^66^, both types of factors would be detected equally. Comparison of our data with factors that occupy their motifs^66^ reveals many such factors display strong DNA binding activity in the ATI assay. The ability to predict DHS positions based on ATI data suggests that much of the nucleosome-competing activity in ES cells is due to TFs that are present in such a high abundance relative to their Kd that they effectively and specifically compete against nucleosome binding. Motif discovery analysis from the DHSs that we could not predict using ATI data also failed to identify motifs that correspond to previously identified pioneer factors that can access nucleosomal DNA^67,68^ (**Supplementary Fig. 8d**), suggesting that such activities either do not rely on complex sequence motifs^68^ or do not occur at so many positions that they would be detectable by motif mining.

Taken together, our findings suggest that cellular transcriptional regulatory network may be much simpler than what has been previously thought, and that the solution to one of the largest remaining problems in the biological sciences – the determination of gene expression from DNA sequence – may be within reach in the near future.

## Methods

### Cell culture and protein extraction

Mouse embryonic stem cells (C57BL/6J; from KCTT center at Karolinska Institutet) were thawed and plated in ES1+LIF medium and cultured in 2i+LIF medium without MEF feeder layers (first set, corresponding to **Fig. 1** and **Fig. 3**) or on low density MEF feeder layers (277,000 irradiated MEFs per 60 mm dish, corresponding to **Fig. 2**) until 70-80% confluence; cells were passaged every two or three days and collected by trypsinization and MEF feeder cells were removed by means of differential adhesion method. The ES1+LIF (250 ml) medium includes: 204 ml knockout DMEM (Gibco, Cat no. 10829-018), 37.5 ml FBS (ES qualified, Sigma, Cat no. F7524), 2.5 ml L-Glutamine (200 mM, Gibco, Cat no. 25030-024), 2.5 ml HEPES (1M, Gibco, Cat no. 15630-056), 2.5 ml Non-Essential Amino Acids (10 mM each, Gibco, Cat no. 11140-035), 0.5 ml β-mercaptoethanol (50 mM, Thermo Scientific, Cat no. 31350-010), 0.25 ml Gentamicin (10 mg/ml, Thermo Scientific, Cat no. 15710-049) and 0.25 ml Leukemia inhibitory factor (1 × 10^6^ U/ml, Millipore, Cat no. ESG1107). The 2i+LIF medium (50 ml) includes: 38.785 ml knockout DMEM, 10 ml KnockOut™ Serum Replacement (Gibco, Cat no. 10828-028), 0.5 ml L-Glutamine (200 mM, Gibco, Cat no. 25030-024), 0.5 ml Non-Essential Amino Acids (10 mM each), 0.1 ml β-mercaptoethanol (50 mM), 50 μl Gentamicin (10 mg/ml), 50 μl LIF (1 × 10^6^ U/ml), 1 μM MEK inhibitor PD0325901 (Miltenyi Biotec, Cat no. 130-103-923), 2 μM GSK-3α/β inhibitor BIO (Sigma, Cat no. B1686). *Drosophila* S2 cells were cultured in Schneider’s *Drosophila* Medium (Thermo Scientific, Cat no. R69007) at 27 °C without CO_2_ and collected by trypsinization. Collected cells were washed once with ice-cold PBS.

For the differentiation of mouse ES cells to different lineages, the ES cells were thawed and plated in ES1+LIF medium and cultured in 2i+LIF medium without feeder layers, and then split to several plates for different treatments. The control ES cells were cultured in 2i+LIF medium without feeder layers; the ES cells for neural differentiation were cultured with 2i medium implemented with 2 μM retinoic acid for 2 days; the ES cells for mesodermal differentiation were first cultured with 2i+LIF medium for 16 hours, and then changed to the mesodermal medium for 30 hours. The mesodermal medium (206 ml) includes: 100 ml IMDM supplemented with GlutaMAX (Thermo Scientific, Cat no. 31980030), 100 ml Ham’s F-12 Nutrient Mix (Thermo Scientific, Cat no. 21765029), 2 ml N2 supplement (100x, Thermo Scientific, Cat no. 17502048), 4 ml B27 supplement (50x, Thermo Scientific, Cat no. 17504044), 0.5 mM ascorbic acid (Sigma, Cat no. A92902), 4.5 × 10^−4^ M monothioglycerol (Sigma, Cat no. M1753), 5 ng/ml VEGF (Thermo Scientific, Cat no. PHC9391), 8 ng/ml Activin A (Thermo Scientific, Cat no. PHG9014) and 0.5 ng/ml BMP4 (Thermo Scientific, Cat no. PHC9534).

All tissues were from one-year old C57BL/6J male mouse. The heart and spleen samples were collected and then cut into small pieces (~ 4 mm) prior to protein extraction; the liver and brain samples were lysed directly without cutting. The nuclear soluble proteins in cells or tissues were extracted by using the “Subcellular Protein Fractionation Kit for Tissues” (Life Technologies, Cat no. 87790). Ice-cold CEB buffer (Life Technologies; 1 ml/100 mg tissue sample or 1 ml/1 million cells) complemented with protease inhibitors (Roche, Cat no. 05892791001) and phosphatase inhibitors (Roche, Cat no. 04906845001) was added, followed by dounce homogenization. Homogenized samples were transferred through a strainer into a clean tube, and then centrifuged at 500 × g for 5 minutes. Subsequently the supernatant was discarded, and ice-cold MEB buffer (Life Technologies) with protease and phosphatase inhibitors was added to extract the membranes of cells, followed by centrifugation at 3000 × g for 5 min. The remaining nuclear pellet was lysed with detergent-free NEB buffer (Life Technologies) with inhibitors, vortexed for 15 seconds, and incubated at 4 °C for 45 minutes with gentle mixing. The supernatant was collected, supplemented with glycerol (5% v/v) and stored at −80 °C in aliquots for future use. All cells were tested regularly for Mycoplasma infection. All tissues were from one-year old C57BL/6J male mouse.

### Lentivirus production and generation of iHeps

The full length ORFs were cloned into pLenti6/V5 lentiviral expression vector using gateway recombination system. Viruses were generated by co-transfection of expression vectors with packaging vectors psPAX2 and pMD2.G (Addgene) into 293FT cells with Lipofectamine 2000 (Thermo Fisher Scientific). The following day the cells were replenished with fresh culture media and virus containing media was collected after 48 h. The virus was concentrated using Lenti-X concentrator (Clonetech).

Human fibroblast cell line CCD-1112Sk was obtained from ATCC (#CRL 2429) and cultured in fibroblast media containing DMEM plus 10% fetal bovine serum with antibiotics. Early passage fibroblasts were seeded on day 0 and transduced on day 1 with cocktails of TFs as previously reported in studies from Morris *et al*.^69^ (Foxa1, Hnf4a, Klf5), Huang *et al*^39^. (Foxa3, Hnf4a, Hnf1a), and Du *et al^38^* (Hnf4a, Hnf1a, Hnf6, Atf5, Prox1, Cebpa), and with the nine specific TFs identified by ATI from mouse liver (Hnf1a, Hnf1b, Dbp, Mafg, Cebpa, Cebpb, Hnf4a, Hnf6/Onecut1, Esrra). The transduction was performed overnight in the presence of 8 μg/ml polybrene in biological duplicates. The virus containing media was replaced the following morning with fresh fibroblast media containing β-mercaptoethanol. On day 3, the cells were changed to defined hepatocyte growth media (HCM, Lonza). On day 7, the cells were replated on type-I collagen coated plates in HCM media in several technical replicates and thereafter the HCM media was changed every second day. On day 29, the cells were passaged to new type-I collagen coated plates and cultured until 6 weeks after transduction.

The cells were harvested from biological duplicates for each condition at several time points for total RNA isolation followed by cDNA synthesis by Transcriptor High Fidelity cDNA synthesis kit (Roche) and real-time PCR using SYBR green (Roche) for primers specific for GAPDH and albumin transcripts. The albumin Ct values were normalized to GAPDH and the mean values of sample replicates were shown for different conditions at indicated time points.

### Active TF identification assay

Protein extract (4 μg) was incubated with 5 μl barcoded double stranded DNA oligos containing 40N random sequences (10 pmol, 900 ng), together with poly-dIdC as a competitor (80 ng) in 1 × binding buffer (140 mM KCl, 5 mM NaCl, 1 mM K_2_HPO_4_, 2 mM MgSO_4_, 100 μM EGTA and 3 μM ZnSO_4_, in 20 mM HEPES, pH = 7.5) at room temprature for 30 min. After incubation, electrophoretic mobility shift assay (EMSA) was carried out on ice for 70 min by using 6% DNA Retardation Gel (Invitrogen, Cat no. EC63652BOX) in 0.5 x TBE buffer (1 mM EDTA in 45 mM Tris-borate, pH 8.0) with 106 V constant voltage. The gel was then dyed with SYBR gold fluorescence dye for 10-20 min and washed with milliQ water. Fragments migrating above the 300 bp marker were collected and eluted in TE buffer (1mM EDTA in 10 mM Tris-Cl, pH 8.0), followed by incubation at 65 °C for 3 h. PCR was carried out using Phusion polymerase (Thermo Sci. Cat no. F530L) to amplify the eluted DNA oligos for 20 cycles with 4 pmol of each primer using Bio-Rad S1000 Thermal Cycler with the following settings: initial denaturation 97 °C for 60 s, denaturation 97 °C for 15 s, annealing 65 °C for 15 s, elongation 72 °C for 40 s, final elongation 72 °C for 180 s. Additional 4 pmol of primers were added before the last cycle of PCR with 20 min elongation time to convert remaining ssDNA into dsDNA. The PCR product was then incubated again with the same extract and the cycle repeated. After 3 to 4 cycles of enrichment, PCR products bearing different barcodes were pooled and purified with QIAquick PCR Purification Kit (Qiagen, Cat no. 28106) for next generation sequencing (NGS) library preparation. NGS was carried out with HiSeq 2000 or 4000 instruments (Illumina). The sequencing data from different cycles were compared with each other to determine the enrichment of specific motifs that relates to the overall DNA binding activities of specific transcription factors.

In principle, ATI analysis using 1 μg of the 40 bp random oligonucleotides (consisting of more than 6 × 10^12^ DNA ligands) can identify exact sequences that are approximately 20 bp long, or redundant sequences that consist of approx. 40 bits of information content. Most known TF motifs are well below this limit, with the exception of long arrays of zinc fingers found in repressor proteins that suppress mobile genetic elements^70,71^. Motifs for some of these proteins cannot be identified by ATI due to lack or extreme rarity of the potentially bound sequences in the initial library pool. To address this, we also run ATI using fragmented mouse genomic sequences.

### ATI assay using genomic fragments

The genomic DNA extracted from the mouse ES cells was sheared to make fragments of approx. 150 bp in size. Then, 270 ng of fragmented genomic DNA was incubated with 5 μg mouse ES cell nuclear extract together with 270 ng poly-dIdC in in 20 μl of 1 × binding buffer. After incubation, electrophoretic mobility shift assay (EMSA) was carried out on ice for 70 min using 6% DNA Retardation Gel (Invitrogen, Cat no. EC63652BOX) in 0.5 × TBE buffer (1 mM EDTA in 45 mM Tris-borate, pH 8.0) with 106 V constant voltage. Fragments above 450 bp (“bound”) and between 150 bp to 300 bp (“unbound”) were collected and prepared for Illumina sequencing using NEBNext Ultra™ DNA Library Prep Kit for Illumina (NEB, Cat no. E7370S). Peak calling was carried out using MACS^72^ program for bound and unbound DNA fragments (setting original sheared DNA as background). The peaks from both samples were separately compared with DHSs from mouse ES cells with BEDOPS program^73^.

### Synthetic DNA library design

The synthetic DNA library design was based on our previous SELEX results^9,20^ that include monomeric and dimeric motifs for TFs. First, a dominating set of motifs, consisting of 921 position weight matrices (PWMs)^9,20^ was extracted from the motifs. Subsequently, the seeds of these motifs were reformatted to include only five different IUPAC nucleotide ambiguity codes, A, T, C, G and N. A set of sequences containing the consensus seed, and seeds with each individual defined base replaced by an N were then generated, leading to the total number of 13,847 sequences. These sequences were then flanked with eight different sets of background sequences in such a way that the total length of all sequences was 35 nucleotides. The background sequences were derived from the human genome that did not contain TF binding sites (based on ChIP-seq data), exons, or high affinity matches to any of the 921 PWMs. Finally, standard Illumina adapters and 3 undefined bases were added to each sequence to generate unique molecular identifiers (UMIs). The DNA library, consisting of single stranded DNA was then amplified by PCR to make double stranded DNA for experiment.

### ATI data analysis

Two methods were utilized to identify the most important transcription factors in different cell types. The *de novo* motif discovery method was based on the “Autoseed” program (see Nitta *et al.*, 2015^13^). In Autoseed, seed sequences representing subsequences whose counts were higher than any other closely related subsequence (using the Huddinge distance metric^13^) were used as seeds. This method is based on direct counts of subsequences (enrichment relative to random sequence) and not on direct comparison between selected and unselected 10-mer sequences, as the latter approach would increase noise due to the low counts of all 10-mer sequences in the unselected library. The method can identify seeds that are separated by a Hamming distance of two or more. Up to 200 highest count local maxima 10 bp sequences (with or without a gap at the center) were used as seeds to generate initial PWM motifs, which were then investigated manually to remove low complexity motifs and motifs that were highly similar. Background correction was performed by using the subtractive method described in Jolma *et al., 2010*^74^. To facilitate comparison of similar motifs, logos were generated in such a way that the frequency of each base was directly proportional to the height of the corresponding letter; because the absolute molecular counts for most motifs range from 10 000 to 100 000, even relatively small differences between motifs are statistically significant; therefore, the motifs are not rescaled based on information content. Counts of motifs were assigned based on number of reads that match the seeds in cycle 4 minus cycle 0. As > 93% of the reads were unique in each case, all reads were used to estimate the absolute number of molecular events. ATI can measure all TF activities, but its sensitivity is limited by the number of observed binding events compared to the background-occurrence of the motif in the random sequence population. In practice, activities that are more than 500-fold lower than the maximal activity identified in a particular cell type are commonly detected.

Known motif enrichment analysis was used to study the enrichment of known motifs. First the number of reads for known motifs were counted by MOODS program^75,76^ before and after enrichment based on a particular cutoff (*p*-value ≤ 0.0001, Score > 11). Subsequently, the enrichment and *p*-value (Winflat^58^) were calculated for each motif; the sensitivity to detect differences using this method was very high, and it could detect highly statistically significant differences whose fold-changes were probably too low to be biologically meaningful. Due to this reason, we reported also the fold-changes in each case.

To analyze the combinatorial binding of TFs in mouse ES cells, seven “strong” motifs (more than 10% of the highest activity as mentioned in **Fig. 2a**) were taken into account. Each read in the sequencing data was analyzed for the presence of perfect matches to each of the seven strong motif seeds and the total number of all the seed matches in each read were counted. For symmetric motifs, only one strand was taken into account; for asymmetric motifs, both strands were analyzed. This analysis revealed that after four cycles, ~5% of reads contained a seed match, and only 0.05% contained matches to more than one seed (**Supplementary Fig. 1b**), indicating that in the early rounds of ATI, the motifs cannot effectively compete against each other. Thus, the presence of only few strongly active motifs in the ATI data cannot be due to over-enrichment of one motif that competes out the other motifs during the enrichment cycles.

### DHS analysis

The DHSs data for different mouse tissues and ES cells were obtained from ENCODE project^50^, including 14 replicates for mouse liver, 7 replicates for mouse brain, 2 replicates for mouse heart, spleen and ES cells. First the BroadPeak data was downloaded, and top 5,000 regions for each replicate were selected based on Signal Values. For tissues with two replicates, the intersected regions were used for downstream analysis, resulting in around 4,000 DHSs for each tissue; for liver, DHSs overlapped by more than 8 replicates were selected to reach similar size of data compared with the other tissues; for brain, DHSs overlapped by more than 4 replicates were selected. For each tissue, the frequencies of all 10-mers were counted in the initial ATI library, and in each ATI enrichment cycle; the fold change for each 10-mer was calculated by comparing the frequencies of it in the last cycle (Cycle4) and first cycle (Cycle1). After that, the DHSs and the 10-mer results of the same tissues were analyzed. First, each DHS region was flanked with adjacent genomic sequences to make a 10 kb region, resulting in ~ 4,000 regions with the length of 10 kb for each tissue and cell type, all these 10 kb regions were then aligned by using the middle of the DHS as center position. A score for each position of the 10 kb sequences was then calculated based on the log2 fold change of the ATI 10-mers. The histogram in **Fig. 4b** was calculated from the average of the scores for all DHSs. For visualization (**Fig. 4b**), the position containing a 10-mer that was ranked at 2000 or higher in enrichment based on ATI was indicated by a blue dot.

The extended 10 kb regions of ES cells were used in the ATI-based prediction of DHSs. All 10-mers enriched in ATI were scored 1, with the remaining scored 0. The scoring of the 10-mers was optimized by trying different cutoffs using a separate training set (setting separately top 0.1%, top 0.5%, top 1%, top 5%, top 10%, top 20%, top 40%, top 60% and top 100% (all) of the 10-mers as score 1 and the remaining 10-mers as score 0, 100% of 10-mers is considered as a negative control). The 10 kb regions were divided to 50 bp bins, and each bin was then assigned with the mean score of all 10-mers inside the bin. The DHS position was then called based on identification of the highest score in a sliding window of 17 bins. The optimal width of the smoothing window was determined by using half of the 10 kb regions as a training set; only the test set data is shown on **Figure 4**.

### Precision-recall analysis

For prediction of DHS regions genome-wide, DHS regions representing the intersection of the two top 30,000 mouse ES cell DHS sequences were selected and for each DHS region, 10 kb control sequences (non-DHS regions) were also taken from both ends of 50 kb windows centered by it. A score was assigned to each 10-mer as the Log2 fold change by comparing the counts of the 10-mer in DHS regions and the control regions derived from chromosomes 12 to 18. For the ATI data, the 10-mer scores were calculated as mentioned in “DHS analysis”. The scores were kept for a fraction of the most enriched 10-mers, with the remaining 10-mers assigned a score of 0. To plot the precision recall curves, a score was assigned to each non-overlapping 1 kb window by adding up the scores of all 10-mers inside the window. Each window was labeled as “DHS” if more than 500 bp of it was covered by a DHS, and “non-DHS” otherwise. Then the precision-recall curve was plotted by predicting the labels of all the windows with their scores using varied thresholds. For the final prediction plots, the fraction of non-zero 10-mer scores was identified by optimizing the area under the curve (AUC) of a precision recall curve using chromosome 11 and 19 DHS data, and the DHS data from the remaining chromosomes was used as ground truth for the prediction.

### Enrichment of DNA-binding proteins using biotinylated ATI ligands

The proteins in the nuclear extract were pulled down by biotinylated DNA as previously reported^77^. First, DNA oligonucleotides were amplified with biotinylated primers (modified with Biotin-TEG) and purified to remove extra primers. Subsequently, 2 μg of biotinylated dsDNA were incubated with 4 μl of high-performance streptavidin Sepharose suspension (GE Healthcare, 17511301) in DNA binding buffer (10 mM HEPES, pH 8.0, 1 M NaCl, 10 mM EDTA, and 0.05% NP40) for 1 h at room temperature by shaking. Beads were then washed twice with DNA binding buffer and twice with protein binding buffer (140 mM KCl, 5 mM NaCl, 1 mM K_2_HPO_4_, 2 mM MgSO_4_, 100 μM EGTA and 3 μM ZnSO_4_, in 20 mM HEPES, pH = 7.5). Nuclear extract from feeder-free mouse ES cells (200 μg in 200 μl supplemented with 2 μg poly-dIdC competitor DNA and EDTA-free complete protease inhibitors (Sigma, Cat no. 000000004693159001) was added to the beads and incubated for 1.5 hours with shaking at room temperature. The beads were then washed with ice-cold low stringency buffer (10 mM Tris-Cl, pH 7.5, 4% glycerol, 500 μM EDTA, 50 mM NaCl) for 10 times and proceeded to on-beads digestion for MS^77^.

### MS sample preparation

For on-beads digestion of captured DNA-binding proteins from the nuclear extract, washed beads were incubated in 50 μl of 25 mM ammonium bicarbonate and 1 mM DTT for 1h in 37 °C. Iodoacetic acid (IAA) was then added to samples to a final concentration of 5 mM and samples were incubated at room temperature in the dark for 10 min. The IAA was then quenched by addition of DTT to a final concentration of 5 mM. Protein samples were then digested, first by using Lys-C protease (0.2 μg/sample, Thermo Scientific, Cat no. 90051) overnight at 37 °C. In the second digestion step trypsin protease (0.1 μg/sample, Thermo Scientific, Cat no. 90057) was added and samples were incubated overnight at 37 °C, and then lyophilized using a speedvac.

For analysis of nuclear proteins, nuclear extracts were prepared as described above and protein concentration was determined (Bio-Rad DC assay). For digestion using filter aided sample prep (FASP), 250 μg of protein sample was mixed with 1mM DTT, 8 M urea, 25 mM HEPES pH 7.6 in a centrifugation filtering unit with a 10 kDa cut-off (Nanosep^®^ Centrifugal Devices with Omega™ Membrane, 10 k). The samples were then centrifuged for 15 min, 14.000g, followed by another addition of the 8 M urea buffer and centrifugation. Proteins were alkylated by 25 mM IAA, in 8 M urea, 25 mM HEPES pH 7.6 for 10 min, centrifuged, followed by two more additions and centrifugations with 4 M urea, 25 mM HEPES pH 7.6. Protein samples were digested on the filter, first by using Lys-C (Thermo Scientific) for 3h in 37 °C, enzyme:protein ratio 1:50. In the second digestion step trypsin (Thermo Scientific), enzyme:protein ratio 1:50 in 50 mM HEPES was added and incubated overnight at 37 °C. After digestion, the filter units were centrifuged for 15 min, 14 000 x g, followed by another centrifugation with 50 uL MilliQ water. Peptides were collected and the peptide concentration determined (Bio-Rad DC assay). For the label-free experiment, peptide samples were cleaned-up individually by solid phase extraction (SPE strata-X-C, Phenomenex) and dried in a SpeedVac.

For the relative quantification (TMT) experiment, peptide samples were pH adjusted using TEAB buffer with pH 8.5 (30 mM final conc.). The resulting peptide mixtures were labelled with isobaric TMT-tags (Thermo Scientific). High labelling efficiency was verified by LC-MS/MS before pooling of samples. Sample clean-up was performed by strong cation exchange solid phase extraction (SPE strata-X-C, Phenomenex). Purified samples were dried in a SpeedVac.

For peptide prefractionation by high resolution isoelectric focusing^42^, 500 μg of the labeled peptide pool was dissolved in 250 μl of rehydration solution (8 M urea, 1% Pharmalyte for pH range 3-10 from GE Healthcare), which was then used to re-swell an immobilized pH gradient (IPG) gel-strip (GE Healthcare) pH 3-10. IEF was then run on an Ettan IPGphor (GE Healthcare) until at least 150 kVh (~1 day running time). After focusing was complete, a well-former with 72 wells was applied onto the strip, and liquid-handling robotics (GE Healthcare prototype) added MilliQ water and, after 3x30 min incubation/transfer cycles, transferred the 72 fractions into a microtiter plate (96 wells, V-bottom, Corning cat. #3894), which was then dried in a SpeedVac.

### Mass spectrometry

Label-free MS of peptides from captured DNA-binding proteins was performed using a hybrid Q-Exactive mass spectrometer (Thermo Scientific). Each sample was resuspended in 10 μl of solvent A (95% water, 5% DMSO, 0.1% formic acid) of which 3 μl was injected. Peptides were trapped on an Acclaim PepMap nanotrap column (C18, 3 μm, 100 Å, 75 μm x 20 mm), and separated on an Acclaim PepMap RSLC column (C18, 2 μm, 100 Å, 75 μm x 50 cm, Thermo Scientific). Peptides were separated using a gradient of A (5% DMSO, 0.1% FA) and B (90% ACN, 5% DMSO, 0.1% FA), ranging from 6 % to 37 % B in 240 min with a flow of 0.25 μl/min. Q Exactive (QE) was operated in a data-dependent manner, performing FTMS survey scans at 70 000 resolution (and mass range 300-1700 m/z) followed by MS/MS (35 000 resolution) of the top 5 ions using higher energy collision dissociation (HCD) at 30% normalized collision energy. Precursors were isolated with a 2 m/z window. Automatic gain control (AGC) targets were 1e^6^ for MS1 and 1e^5^ for MS2. Maximum injection times were 100 ms for MS1 and 150 ms for MS2. The entire duty cycle lasted ~1s. Dynamic exclusion was used with 60 s duration. Precursors with unassigned charge state or charge state 1 were excluded. An underfill ratio of 1% was used.

LC-MS of TMT labeled peptides from nuclear extracts was also performed using a hybrid Q-Exactive mass spectrometer (Thermo Scientific). For each LC-MS/MS run, the auto sampler (Dionex UltiMate™ 3000 RSLCnano System) dispensed 15 μl of solvent A (95% water, 5% DMSO, 0.1% formic acid) to the well in the 96 well plate, mixed, and 7 μl proceeded to injection. Peptides were trapped on an Acclaim PepMap nanotrap column (C18, 3 μm, 100 Å, 75 μm x 20 mm), and separated on an Acclaim PepMap RSLC column (C18, 2 μm, 100Å, 75 μm x 50 cm, Thermo Scientific). Peptides were separated using a gradient of A (5% DMSO, 0.1% FA) and B (90% ACN, 5% DMSO, 0.1% FA), ranging from 6 % to 37 % B in 50 min with a flow of 0.25 μl/min. QE was operated as described above.

Label-free MS of proteins in the nuclear extract was performed using Orbitrap Fusion™ Tribrid mass spectrometer (Thermo Scientific). Before the analysis, peptides were separated using an Ultimate 3000 RSLCnano system. Samples were trapped on an Acclaim PepMap nanotrap column (C18, 3 μm, 100Å, 75 μm × 20 mm), and separated on an Acclaim PepMap RSLC column (C18, 2 μm, 100Å, 75 μm × 50 cm), (Thermo Scientific). Peptides were separated using a gradient of A (5% DMSO, 0.1% FA) and B (90% ACN, 5% DMSO, 0.1% FA), ranging from 6 % to 37 % B in 240 min with a flow of 0.25 μl/min. The Orbitrap Fusion was operated in a data dependent manner, selecting top 10 precursors for sequential fragmentation by HCD and CID. The survey scan was performed in the orbitrap at 120,000 resolution from 350-1550 m/z, with a max injection time of 50 ms and target of 2 × 10^5^ ions. Precursors were isolated by the quadrupole with a 1.4 m/z window and a 0.5 m/z offset, and put on the exclusion list for 30s. Charge states between 2 and 7 were considered for precursor selection. For generation of HCD fragmentation spectra, a max ion injection time of 100 ms and AGC target of 1 × 10^5^ were used before fragmentation at 37% normalized collision energy, and analysis in the orbitrap at 30,000 resolution. For generation of CID fragmentation spectra, a max ion injection time of 100 ms and AGC target of 1 × 10^4^ were used before fragmentation at 35% activation energy, activation Q 0.25, and analysis in the iontrap, using normal scan range and rapid scan rate.

### Peptide and protein identification

For the label free capture experiment, MS raw files were searched using Sequest-percolator under the software platform Proteome Discoverer 1.4 (Thermo Scientific) against Uniprot mouse database (version 2016_10,canonical and isoforms, 85832 protein entries) and filtered to a 1% FDR cut off (peptide spectrum match level). A maximum of 2 missed cleavages were used together with: carbamidomethylation (C) set as fixed modification, and oxidation (M) as variable modification. We used a precursor ion mass tolerance of 10 ppm, and a product ion mass tolerance of 0.02 Da for HCD spectra. For calculation of precursor ion area, a mass precision of 2 ppm between scans was used, and the average area of the top 3 PSMs for each protein group was used to calculate protein area. Only unique peptides in the data set were used for quantification. In total the database search resulted in the identification of 3889 proteins (**Supplementary Table 3**). TFs were assigned to related motifs detected in ATI based on the current database (HT-SELEX motifs, CIS-BP, TOMTOM^9,13,40,41^). If only one TF was assigned to a motif, this particular TF was regarded as the candidate TFs; if more than one TF were assigned to a motif, TFs with more than 20% of the highest abundance (based on values in “Aver_area(c4)”) were selected as the candidate TFs which are dominant in the cells (**Supplementary Table 1**). These parameter values should be considered only an example, as the optimal cut-offs depend on the TFs and purpose of the individual projects.

For the nuclear extracts analysis, MS raw files were searched using Sequest-percolator under the software platform Proteome Discoverer 1.4 (Thermo Scientific) against Uniprot mouse reference database (version 2014_03, canonical only, 43386 protein entries) and filtered to a 1% FDR cut off (peptide spectrum match level). For TMT experiments, a maximum of 2 missed cleavages were used together with: carbamidomethylation (C), TMT/ 10-plex (K, N-term) set as fixed modifications, and oxidation (M), as variable modification. We used a precursor ion mass tolerance of 10 ppm, and a product ion mass tolerance of 0.02 Da for HCD spectra. Quantification of reporter ions was done by Proteome Discoverer on HCD-FTMS tandem mass spectra using an integration window tolerance of 10ppm. Only unique peptides in the data set were used for quantification. In total the database search resulted in the identification and quantification of 8578 proteins in the TMT experiment (**Supplementary Table 9**).

For label free analysis, a maximum of 1 missed cleavage were used together with: carbamidomethylation (C), set as fixed modifications, and oxidation (M), as variable. We used a precursor ion mass tolerance of 12 ppm, and a product ion mass tolerance of 0.02 Da for HCD spectra and 0.36 for CID spectra. For calculation of precursor ion area, a mass precision of 3 ppm between scans were used, and the average area of the top 3 PSMs for each protein group were used to calculate protein area. In total the database search resulted in the identification of 6239 proteins in the label free experiment (**Supplementary Table 10**).

## Acknowledgments

We thank Drs. Jian Yan, Eevi Kaasinen, Bernhard Schmierer, Yimeng Yin for critical review of the manuscript, and Sandra Augsten, Lijuan Hu and Poomy Pandey for technical assistance. This work was supported by Center for Innovative Medicine at Karolinska Institutet, Knut and Alice Wallenberg Foundation, Göran Gustafsson Foundation and Vetenskapsrådet.

## Author Contributions

I.K.S. collected mouse tissue samples; B.W. extracted proteins, performed ATI experiments and analyzed the data; Fan.Z. and Fangjie.Z. did the DHS analysis; B.S. did the iHep reprogramming experiment; L.M.O. and J.L. did the mass spectrometry experiments and data analysis; A.J., T.K. and M.T. helped to supervise the project or related experiments. B.W. and J.T. wrote the manuscript.

## Author Information

All next generation sequencing data have been deposited to European Nucleotide Archive (ENA) under Accession PRJEB15639. All computer programs and scripts used are either published or available upon request. The authors declare no competing financial interests. Requests for materials should be addressed to J.T.

## Supplementary Figures

**Supplementary Figure 1.**
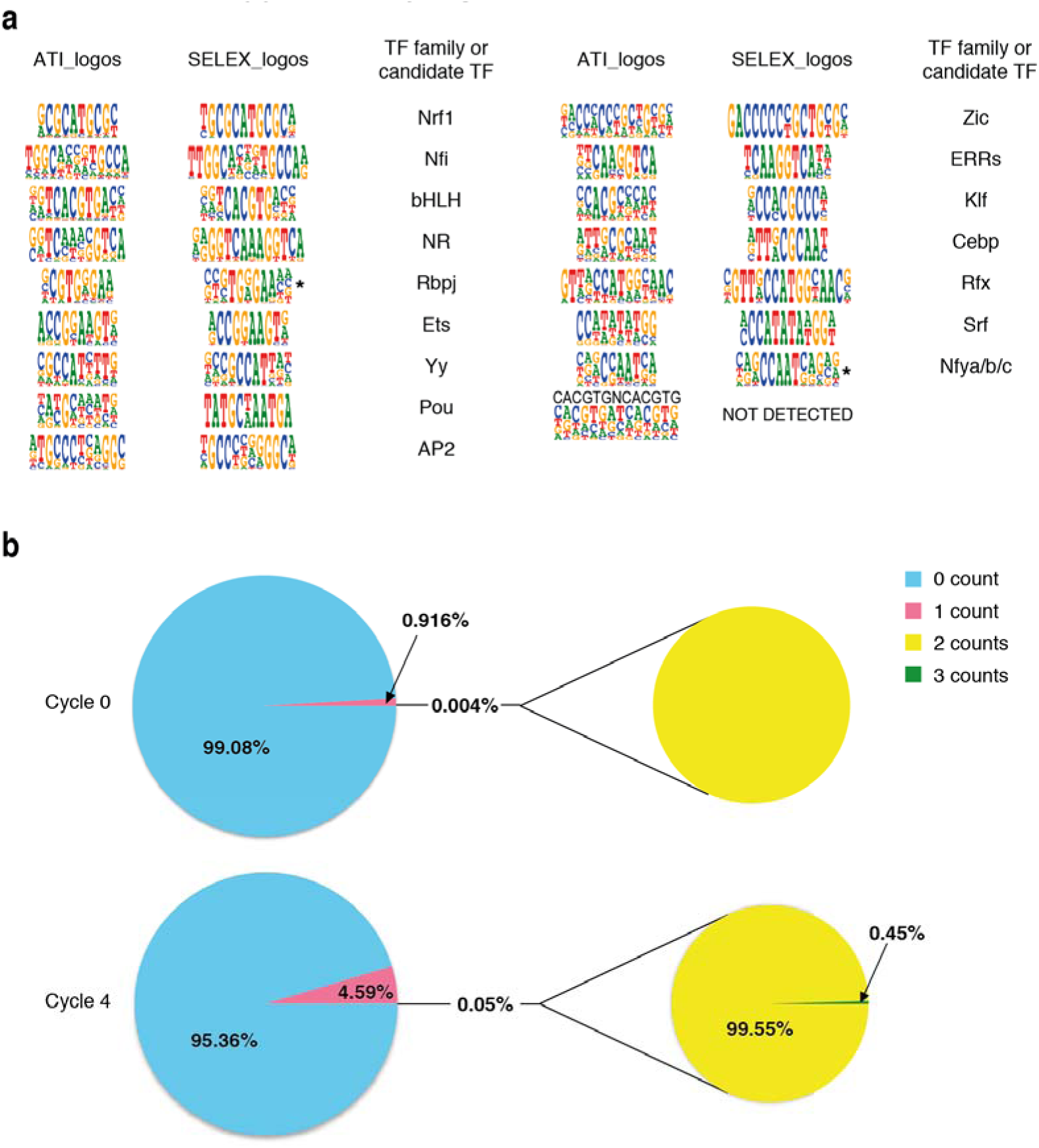
Analysis of motifs detected in ATI assay using extract from feeder free mouse ES cells. **a,** The motifs detected in ATI assay by using the mouse ES cells extract are compared with reference motifs detected by using bacterial expressed pure proteins in HT-SELEX^9^. The reference binding motif of TF Rbpj (asterisk) is from T. Tun *et al.*^21^; the binding motif of TF Nfy (asterisk) is from the HOCOMOCO database^78^. The TF families or specific TFs are proposed based on the comparison of the motifs. **b,** The pie charts indicate the percentage of reads containing different numbers (counts) of seed matches to the strong motifs found in mouse ES cells. Top: matches in the original input DNA pool (“Cycle 0”). Bottom: matches in the ATI-enriched DNA pool (“Cycle 4”).

**Supplementary Figure 2.**
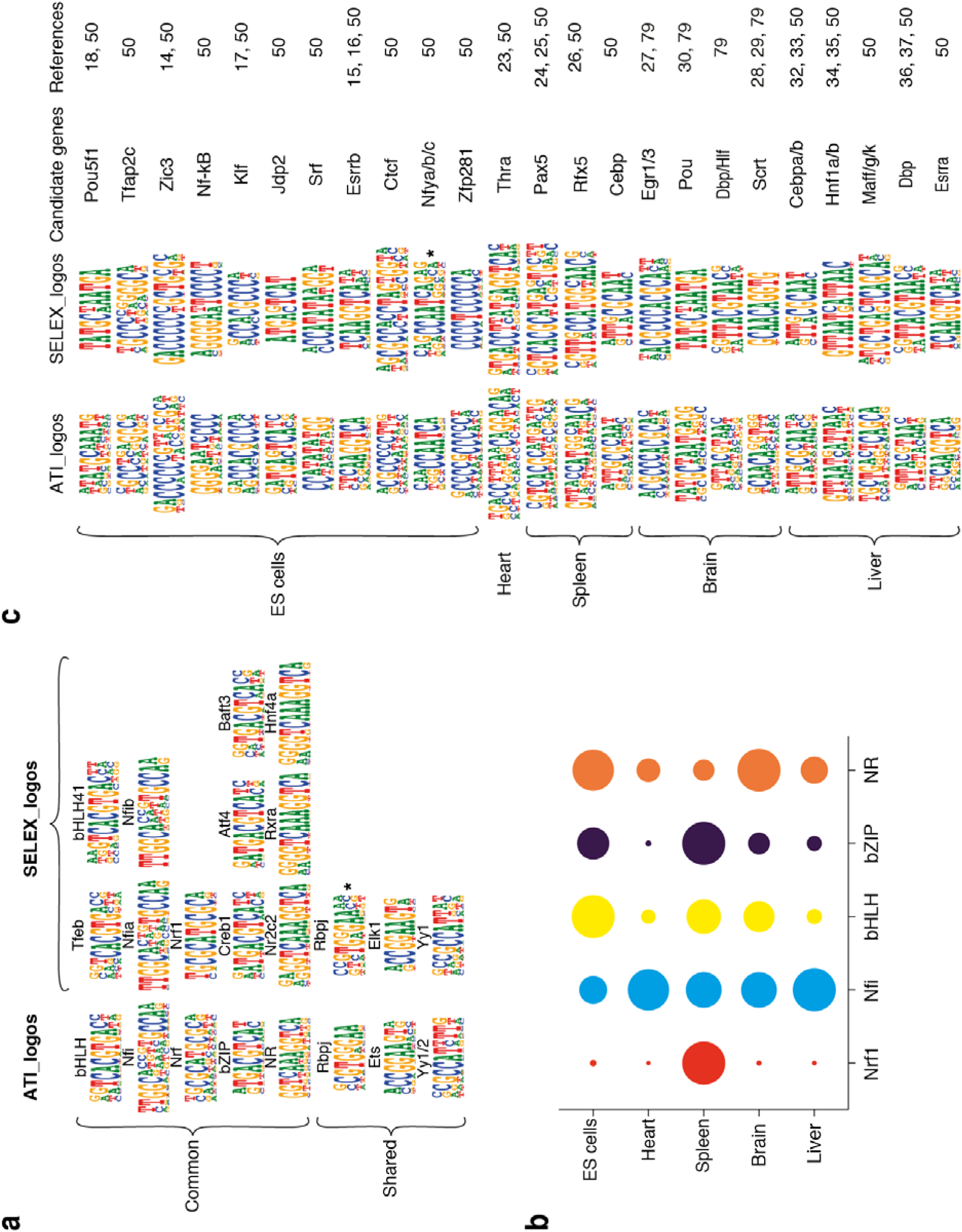
Motif analysis of ATI data using nuclear extract from different types of mouse tissues and cell lines. **a,** “Common” and “Shared” motifs that are found in different mouse cell and tissue types are compared with corresponding motifs detected by using bacterially expressed pure proteins using HT-SELEX^9^. There is one exception that is not in the SELEX database and corresponds to motif of TF Rbpj; the reference motif of Rbpj (asterisk) is from T. Tun *et al.*^21^, 1994. **b,** The activities of all five “common” TFs detected in all tested samples are compared based on the absolute molecular counts^12^ of each motif in the sequencing data. The areas of circles indicate the activities of the five common motifs in the indicated tissues. Data from the last cycle (cycle 4) are used as signals, and data from the previous cycle (cycle 3) are used as background to determine enrichment in one single ATI cycle. For each motif, the activities are normalized by setting its highest activity in any of the tissues as 1. **c,** The motifs detected in ATI assay by using different mouse cell and tissue samples are compared with the similar motifs detected by using bacterial expressed pure proteins in HT-SELEX^9^. The binding motif of TF Nfy (asterisk) is from the HOCOMOCO database^78^. For TFs having the unique binding motifs, the results are validated by the mRNA expression of those TFs; for TFs sharing the same binding motifs, the specific members are proposed based on the mRNA expression levels^50,79^ and functional data from previous studies.

**Supplementary Figure 3.**
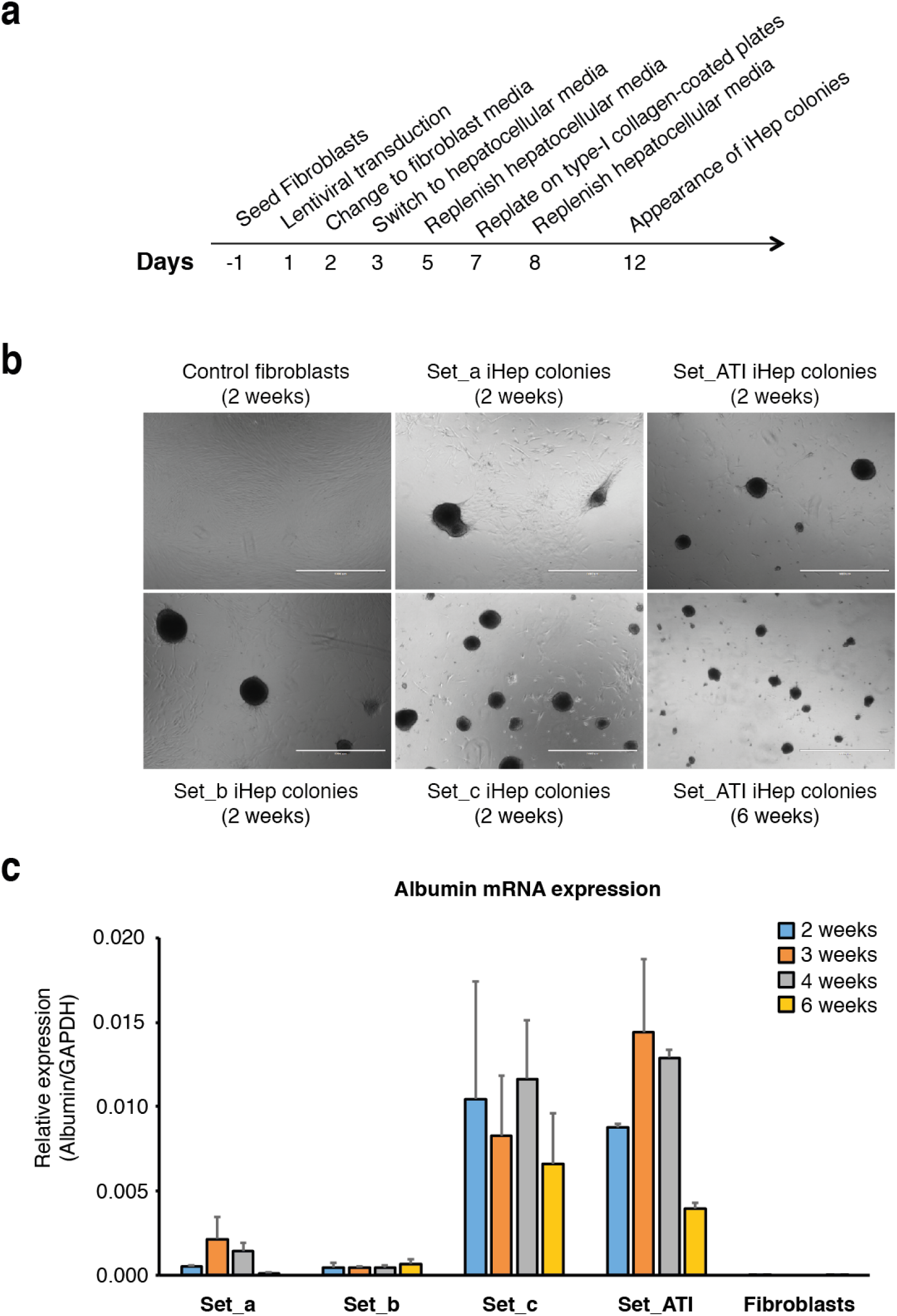
Reprogramming of human induced hepatocytes confirms the results of ATI assay. **a,** Reprogramming timeline for direct conversion of human fibroblasts to induced hepatocytes (iHep). **b,** Bright field images of iHep colonies from human fibroblasts after lentiviral transduction of transcription factors (TF) combinations previously reported in Morris *et al*.^69^ (Set_a; Foxa1, Hnf4a, Klf5), Du *et al*.^38^ (Set_b; Hnf4a, Hnf1a, Hnf6, Atf5, Prox1, Cebpa), Huang *et al*.^39^ (Set_c; Foxa3, Hnf4a, Hnf1a) and factors identified by ATI in mouse liver (Set_ATI; Hnf1a, Hnf1b, Dbp, Mafg, Cebpa, Cebpb, Hnf4a, Hnf6, Esrra). **c,** Expression levels of the liver-specific marker gene Albumin in iHep cells normalized to GAPDH levels by qRT-PCR from two biological replicates using previously reported TF cocktails and ATI-identified TF combinations. Bars indicate standard deviation of biological replicates (n=2).

**Supplementary Figure 4.**
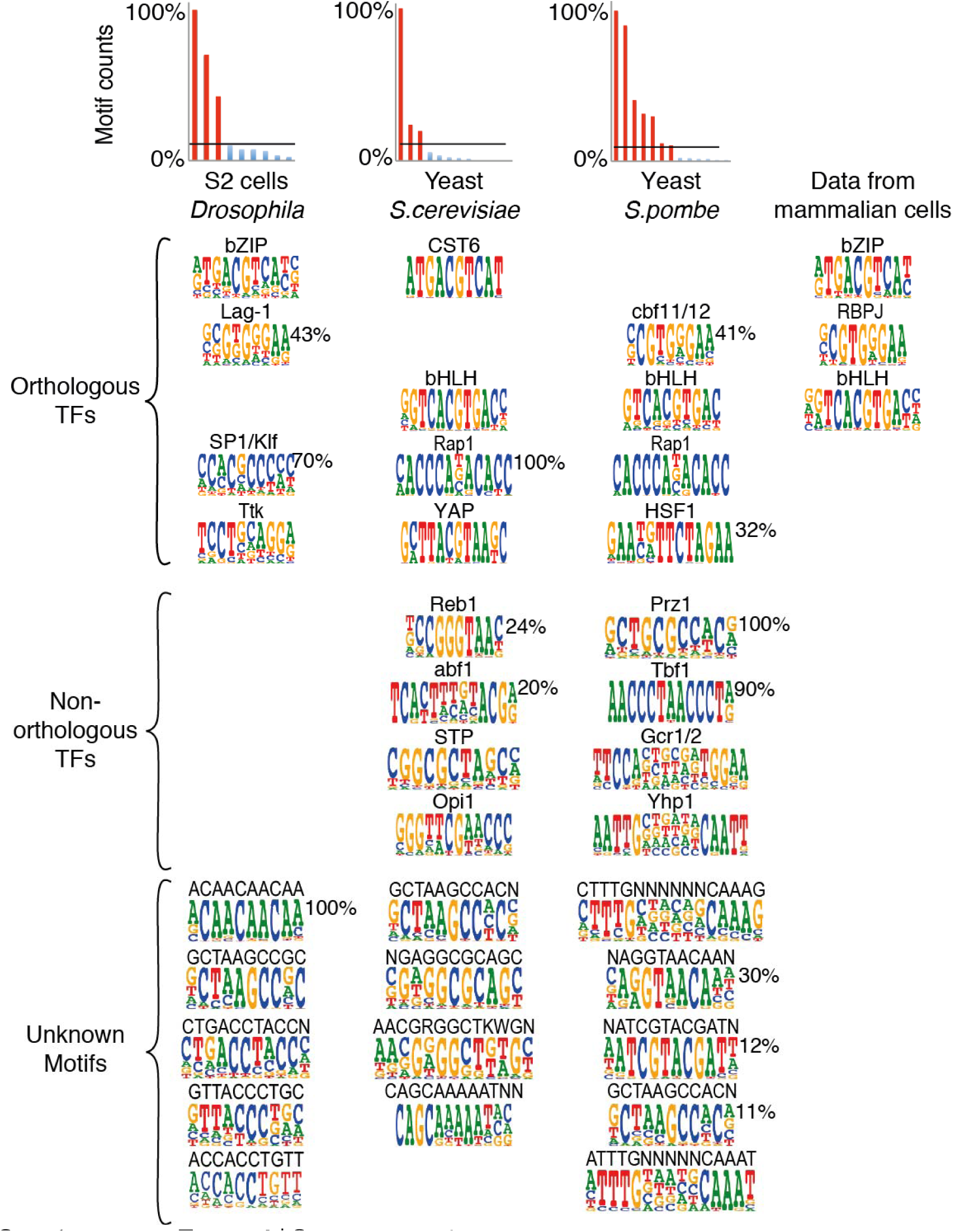
ATI assay from multiple species reveals deep conservation of TF activity. ATI analysis of four different species indicates that the assay can identify TF activities from a wide variety of organisms. The names of the TF that bind to motifs that are similar to those identified using ATI are shown above the sequence logos. Histograms on top show background corrected absolute molecular counts^12^ (y-axis, Motif counts) of all discovered motifs at enriched ATI cycle; for each sample the highest count is normalized to 100%. Counts more than 10% of the maximum are indicated by red bars; the relative activities of them are shown on the right corner of the corresponding sequence logos. Note that many TFs have similar specificities between the species, and that out of the six motifs that are active in most mouse tissues, two (Rbpj/Cbf11 and Tfe/Cbf1) are also highly active in the yeast *S.pombe*, and two (CST6/Creb and Tfe/Cbf1) are highly active in *S.cerevisiae*. Note also that several motifs that could not be assigned to a known TF based on the literature were detected in the species using the indicated seed sequences.

**Supplementary Figure 5.**
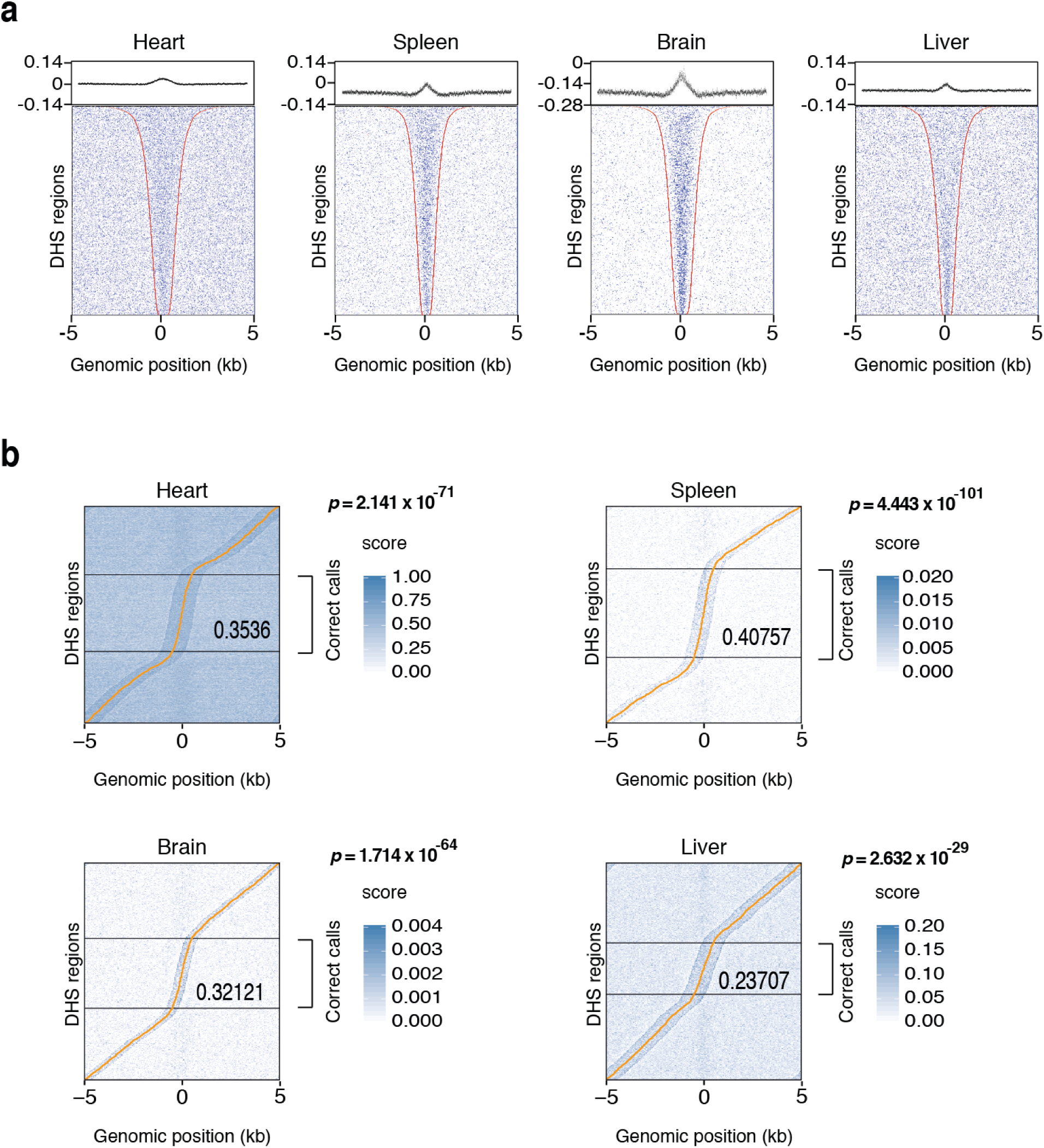
Enrichment of ATI 10-mers in DNase I hypersensitive sites from different mouse tissues. **a,** ATI enriched 10-mers from mouse tissues are also enriched in DNase I hypersensitive sites from the corresponding tissues. The dot plots show matches to enriched ATI 10-mers in DNase I hypersensitive sites from the indicated mouse tissues. In each dot plot, each row indicates one DHS region from the relative mouse tissues that is flanked with its genomic sequences. Red dots indicate the boundaries of the DHS regions, blue dots indicate positions of top 2000 ATI-enriched 10-mers out of all 4^10^ (~ 1 million) 10-mers. The graph on top shows the average of scores for each 10-mer at each position across the rows. **b,** Prediction of DHS regions by using the 10-mer data from the ATI assay. DHSs are sorted by position of the prediction call (yellow line). Black horizontal lines separate accurate DHS calls (middle) from calls more than 500 bp off the known DHS center that is located at the x-axis position 0 in all cases. The fraction of predictions within ± 500 bp of the center and the corresponding p-value for null model where position calls are randomly distributed are also indicated. For each tissue, the scoring of the 10-mers was optimized by trying different cutoffs using a separate training set (setting separately top 0.1%, top 0.5%, top 1%, top 5%, top 10%, top 20%, top 40%, top 60% and top 100% of the 10-mers as score 1 and the remaining 10-mers as score 0, top 100% 10-mers is considered as negative control).

**Supplementary Figure 6.**
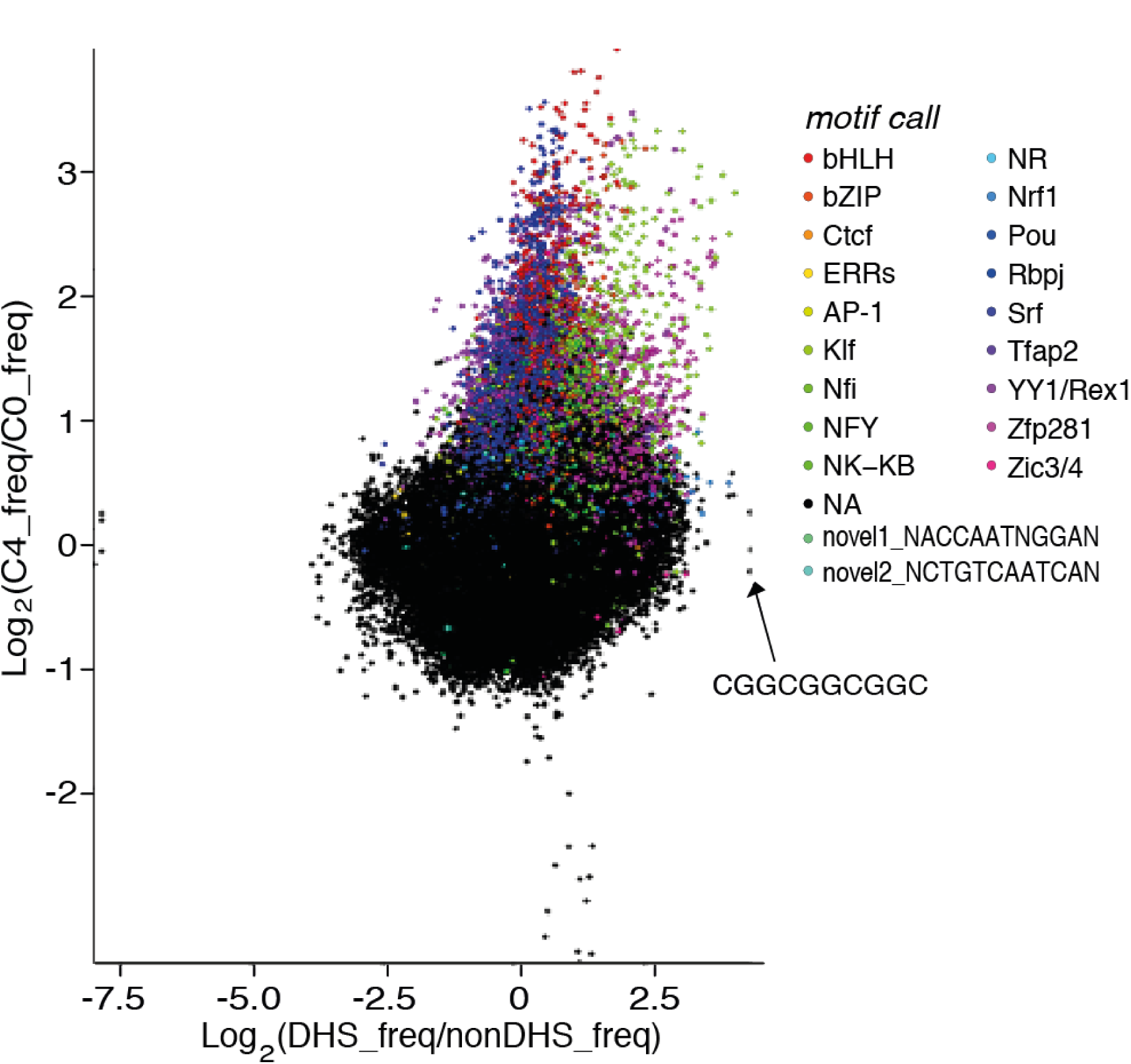
Comparison of subsequences enriched in ATI and in DHS regions from mouse ES cells. **b,** The enrichment of all 10-mer sequences in the ATI data (y-axis) and DHS data (x-axis) from ES cells is shown. X-axis indicates the log2 fold change of 10-mer counts in DHS regions compared with non-DHS regions; y-axis indicates fold change of 10-mer counts in ATI enriched DNA pool (Cycle 4) compared with original pool (Cycle 0). Coloring of the dots indicates 10-mers that are similar to the motifs shown on the right; black dots indicate the 10-mers that are not similar to any motifs. One such 10-mer sequence (“CGGCGGCGGC”) is shown as an example; it displays high enrichment in DHS regions but no enrichment in ATI.

**Supplementary Figure 7.**
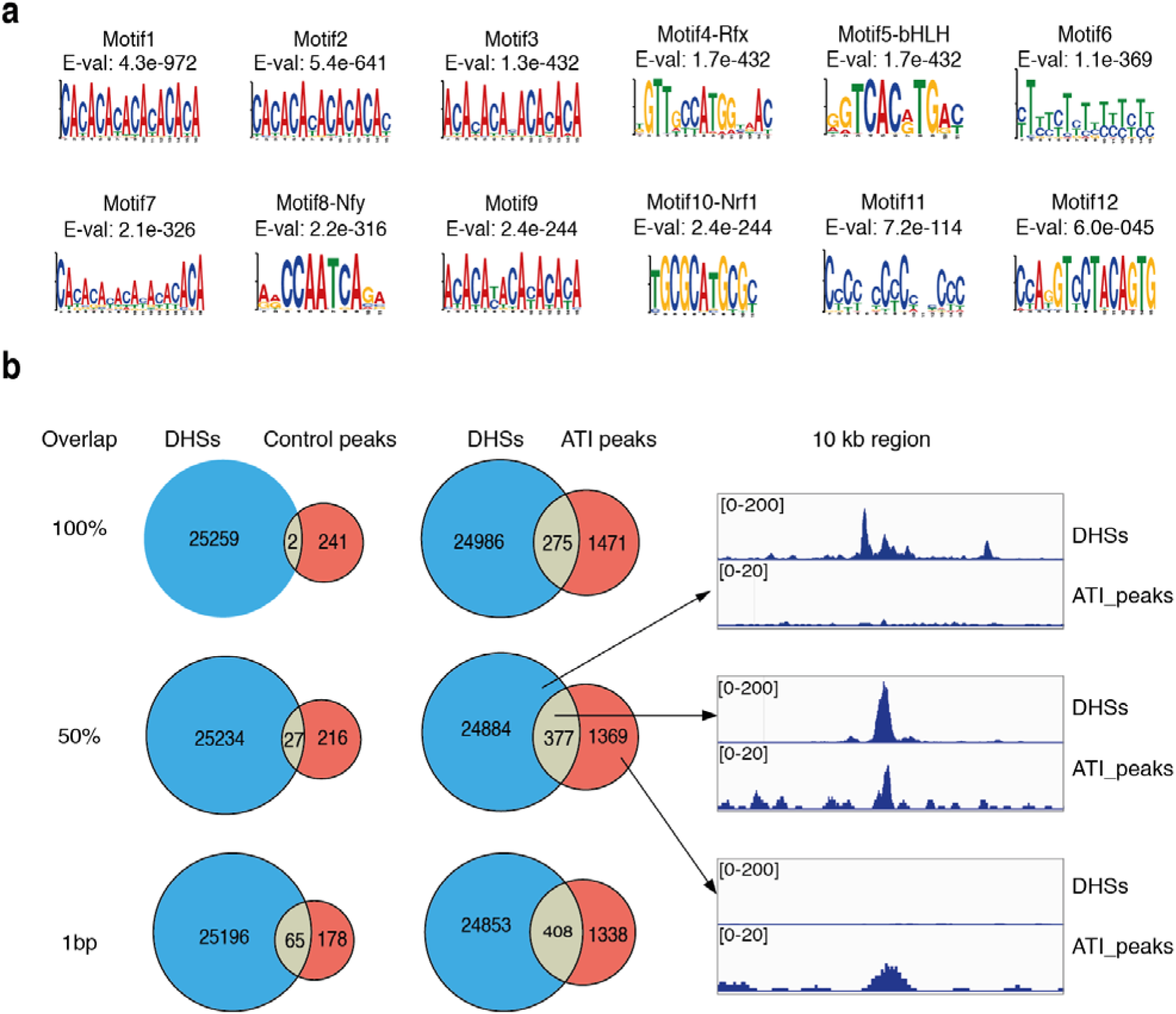
ATI assay by using genomic DNA in mouse ES cells. **a,** *De novo* motif mining of genomic fragments bound by nuclear extract from mouse ES cells. Top twelve motifs are shown with the corresponding E-values. Motifs 4, 5, 8 and 10 were similar to motifs detected in the ATI assay. **b,** Overlap between the 25,261 DHS regions from mES cells (DHSs) and the peaks called from genomic fragments bound by nuclear extract (ATI peaks) or the peaks called from genomic fragments not bound by nuclear extract (Control peaks). The peak analyzed is considered overlapping with DHSs if not less than the indicated percentage or length of the peak overlaps with the DHS regions. The numbers in corresponding areas indicate numbers of DHSs or peaks. The right panel shows three specific loci exemplifying the non-overlapped DHS regions without ATI peaks (top), DHS and ATI overlapped regions (middle) and non-overlapped ATI peaks (bottom).

**Supplementary Figure 8.**
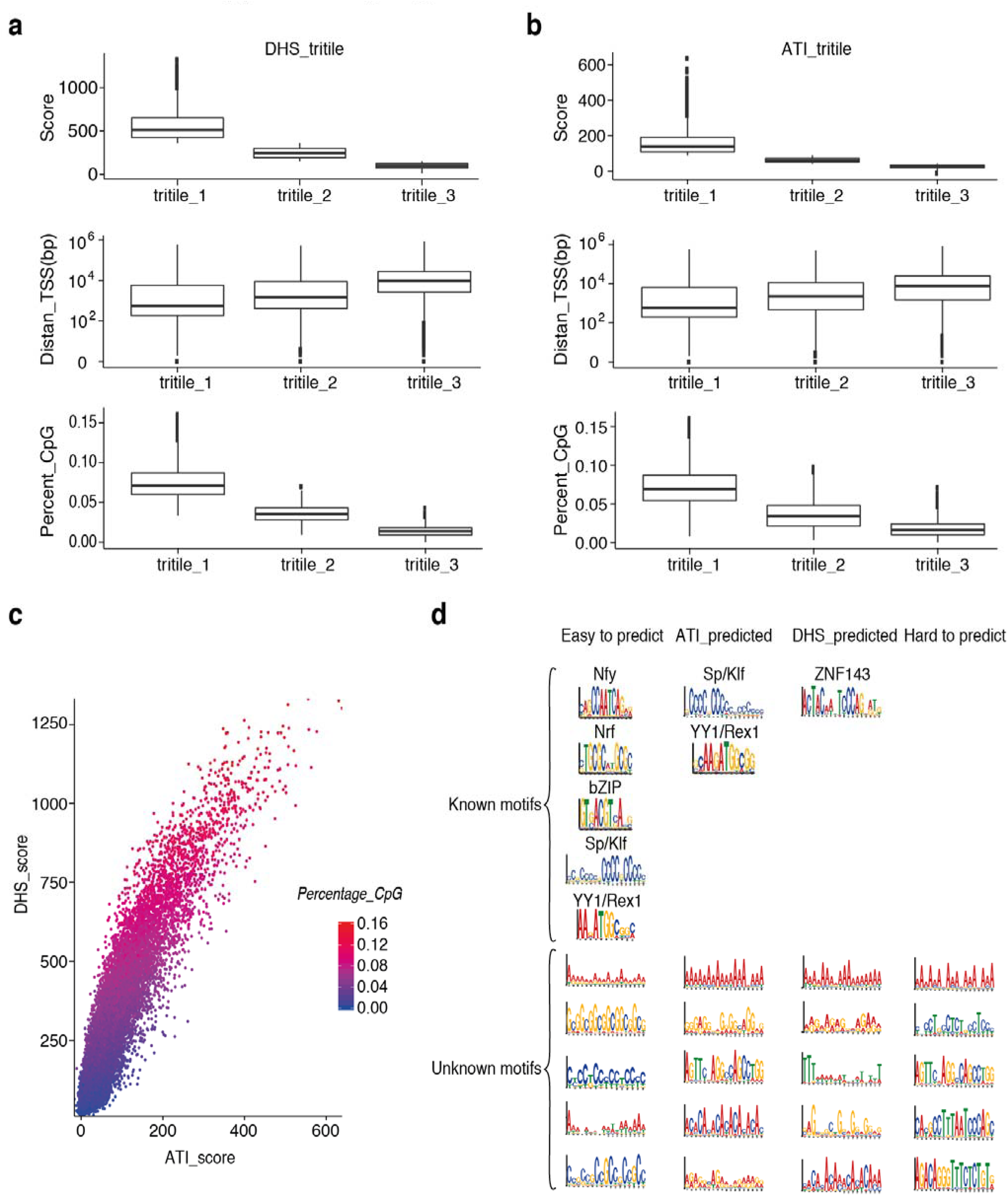
Analysis of features of different categories of DHS fragments. **a-b,** Comparison of features between DHS regions that are easy to predict (tritile_1), intermediate (tritile_2) and hard to predict (tritile_3) using DHS (**a**) or ATI (**b**) 10-mers. The DHS fragments are 1 kb non-overlapping “DHS” fragments used for the final prediction in the Precision-recall analysis. Top: prediction score (sum of the scores of all 10-mers inside the window). Middle: distance from TSS. Bottom: percentage of CpG dinucleotides. Note that hard to predict DHSs using ATI and DHS 10-mer data tend to be farther from a TSS and have a low CpG content. **c,** The correlation between the ATI and DHS total prediction scores for all the 1 kb DHS fragments used for the final prediction in the Precision-recall analysis. Each dot represents one fragment and the color indicates the percentage of CpG dinucleotides within the fragment. **d,** *De novo* motif mining of four different types of DHS regions from mouse ES cells. The different types of DHS regions are generated from intersection of different categories of DHS fragments: easy to be predicted with both ATI and DHS 10-mer data (intersection of tritile_1 of DHS-based prediction in **a** and tritile_1 of ATI-based prediction in **b**, “Easy to predict”), easy to be predicted with ATI 10-mer data but hard to be predicted with DHS data (intersection of tritile_3 of DHS-based prediction in **a** and tritile_1 of ATI-based prediction in **b**, “ATI_predicted”), easy to be predicted with DHS 10-mer data hard to be predicted with ATI data (intersection of tritile_1 of DHS-based prediction in **a** and tritile_3 of ATI-based prediction in **b**, “DHS_predicted”) and hard to be predicted with both types of data (intersection of tritile_3 of DHS-based prediction in **a** and tritile_3 of ATI-based prediction in b, “Hard to predict”). The “known motifs” indicate the motifs can be assigned to the known motifs based on current knowledge. All known motifs with E-value less than 0.01, and top five unknown/repetitive motifs are shown.

**Supplementary Table 1.**
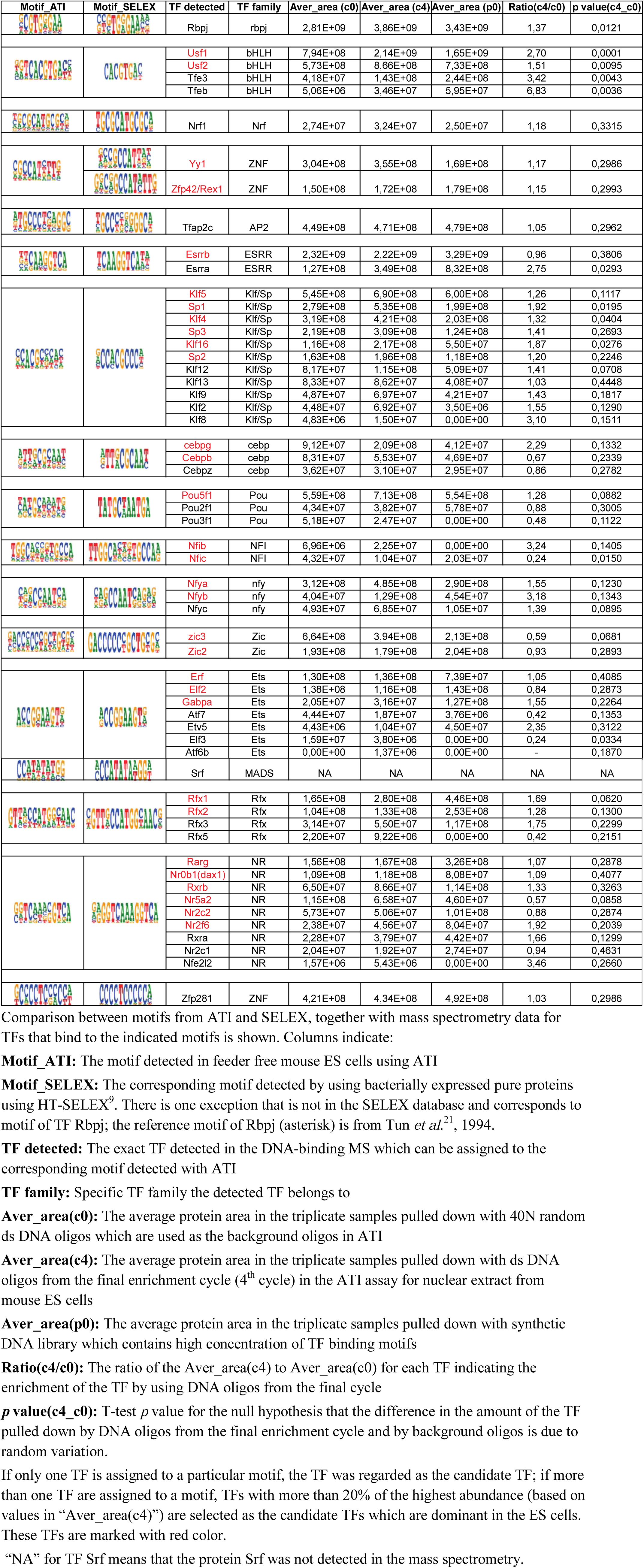
Comparison of the motif analysis result and MS identification result in ATI from nuclear extract of mouse ES cells

